# tRNA Methylation Resolves Codon Usage Bias at the Limit of Cell Viability

**DOI:** 10.1101/2022.05.14.491930

**Authors:** Isao Masuda, Yuka Yamaki, Rajesh Detroja, Somnath Tagore, Henry Moore, Sunita Maharjan, Yuko Nakano, Thomas Christian, Ryuma Matsubara, Todd Lowe, Milana Frenkel-Morgenstern, Ya-Ming Hou

## Abstract

The codon usage of each genome is closely correlated with the abundance of tRNA isoacceptors. How codon-usage bias is resolved by tRNA post-transcriptional modifications is largely unknown. Here we demonstrate that the *N*^1^-methylation of guanosine at position 37 (m^1^G37) on the 3’-side of the anticodon, while not directly responsible for reading of codons, is a neutralizer that resolves differential decoding of proline codons. A genome-wide suppressor screen of a non-viable *E. coli* strain, m^1^G37-lacking, identifies *proS* suppressor mutations, indicating a coupling of methylation with tRNA prolyl-aminoacylation that sets the limit of cell viability. Using these suppressors, where prolyl-aminoacylation is decoupled from tRNA methylation, we show that m^1^G37 neutralizes differential translation of proline codons by the major isoacceptor. Lacking of m^1^G37 inactivates this neutralization and exposes the need for a minor isoacceptor for cell viability. This work has medical implications for bacterial genomes that exclusively use the major isoacceptor for survival.

## Introduction

Codon usage is a unique feature of each gene and each genome and it impacts the fitness of each cell. In the degeneracy of the genetic code, proteins can be coded in multiple ways using different sets of synonymous codons, which are not translated equally in speed or quality. Each codon choice makes a specific demand for the supply of the tRNA isoacceptors that have the matching anticodons. This supply-to-demand balance is broadly observed throughout the domains of life (e.g., *Escherichia coli* (Ikemura, 1981), *Saccharomyces cerevisiae* (dos Reis et al., 2004), *Caenorhabditis elegans* (Duret, 2000), *Drosophila spp*. (Moriyama and Powell, 1997), and eukaryotes in general (Sabi and Tuller, 2014)). It ensures that frequent codons are translated by abundant tRNAs to speed up local translation, whereas rare codons are met with rare tRNAs to slow down local translation (Hanson and Coller, 2018). The impact of this balance is manifested in multiple aspects of a cellular life, including expression level of protein, accuracy of translation, maturity of protein folding, and stability of mRNA (Rak et al., 2018). However, the quality of a codon-anticodon pair is determined not only by the abundance of each isoacceptor, but also by the epigenetic modifications to the anticodon of the isoacceptor that are synthesized post-transcriptionally (de Crecy-Lagard and Jaroch, 2021; Maraia and Arimbasseri, 2017; Pan, 2018). The evolution of epigenetic modifications is an important mechanism to explore the wobble rule of the genetic code, where the wobble nucleotide of the anticodon (at position 34 of tRNA) can form non-canonical base pairing interactions with the 3^rd^ nucleotide of a codon, thus enabling one isoacceptor to translate multiple codons (Agris et al., 2018). Indeed, many genomes, across all three domains of life, encode just one isoacceptor for all synonymous codons. How a single isoacceptor translates all synonymous codons with the speed and quality that is necessary to support life is poorly understood.

The wobble nucleotide at position 34 of tRNA hosts one of the highest chemical diversity and density of post-transcriptional modifications. At this position, several modifications have the ability to expand the anticodon-codon pairing from a canonical W-C structure to wobble structures, thus impacting translation across multiple families of tRNAs. For example, the position-34 modification of adenosine-to-inosine (A34-to-I34), expanding base pairing from A34 with U to I34 with C, U, and A (Torres et al., 2014), is conserved in Eukarya in eight families of tRNAs (Ala, Arg, Ile, Leu, Pro. Ser, Thr, and Val). This expansion is important for neuron health in human (Ramos et al., 2020), and is most critical in organisms that use NNC as the preferred codons, but lack the corresponding tRNA(GNN) isoacceptors, implying that the I34-modified tRNA(ANN) isoacceptors are solely responsible for translation of NNC codons. Indeed, I34-knockout (KO) is lethal to cells, while I34-deficiency (KD) changes the profile of tRNA distribution and re-programs expression of NNC-enriched mRNAs (Lyu et al., 2020). Similarly, several other position-34 modifications in tRNA are also known to expand the capacity of decoding (Agris *et al*., 2018). Deficiency of these modifications also produces changes of gene expression (Begley et al., 2007; Chionh et al., 2016; Nedialkova and Leidel, 2015; Zinshteyn and Gilbert, 2013). However, not all wobble base pairings are created equal, raising the question of whether some wobble base pairings may be deficient in thermodynamic stability and thus would be discriminated against during the high demand of global protein synthesis to support life. This question is most relevant to the deficiency that threatens cell viability, which impacts on the genome structure in evolution.

We hypothesize that the potential deficiency of a wobble base pairing can be compensated by additional post-transcriptional modifications in tRNA. Of interest is the position-37 modification on the 3’-side of the anticodon, which is similarly populated in the density and diversity of chemical structures as the position-34 modification. Importantly, while the position-37 modification is not within the anticodon triplet and is not directly involved in the anticodon-codon pairing, it is often conserved with the position-34 modification or other anticodon-loop modifications in a so-called “modification circuit” (Han and Phizicky, 2018). For example, the t^6^A37 (*N*^6^-threonyl-carbamoyl adenosine) modification in *E. coli* Leu(CAA) and Leu(UAA) is in a circuit with the 2’-*O*-methylation of C34/U34 (Cm/Um); the yW37 (wybutosine) modification in yeast and human tRNA^Phe^ is in a circuit with Cm32 and Gm34; the t^6^A37 modification in *S. cerevisiae* tRNA^Thr^ is in a circuit with m^3^C32; and the i^6^A37 (*N*^6^-isopentenyl adenosine) modification in *S. pombe* and *S. cerevisiae* is in a circuit with m^3^C32 of tRNA^Ser^. These circuits are thought to enhance enzyme modification specificity, such that the first modification in the circuit is a recognition element that facilitates the second modification (Han and Phizicky, 2018). While this notion rationalizes the ordered enzymatic synthesis of the modification circuit, which is supported by some evidence (Han and Phizicky, 2018; Masuda et al., 2019), the biological question of why the circuit is formed is unanswered. Particularly, whether the position-37 modification compensates the deficiency of the position-34 modification to improve the quality of anticodon-codon pairing at the level necessary for cell viability is unknown.

Here we address this question focusing on the m^1^G37 methylation that is invariably associated with tRNAs of the Pro, Leu, and Arg families, and is specifically associated with isoacceptors of Pro(CGG, GGG, and UGG), Leu(CAG, GAG, and UAG), and Arg(CCG) (Bjork and Hagervall, 2014). Deficiency of m^1^G37 stalls ribosomes at most of the associated codons (Masuda et al., 2021), clearly indicating a role of the methylation in regulating the quality of anticodon-codon pairing. However, the biology of m^1^G37 is complex. It is required for accuracy of the translational reading frame (Gamper et al., 2021a; Gamper et al., 2021b; Gamper et al., 2015; Hoffer et al., 2020); loss of m^1^G37 results in +1 frameshifting and premature termination of protein synthesis, leading to cell death (Bjork et al., 1989; Masuda *et al*., 2019). It is also required for efficient and specific aminoacylation of tRNA (Masuda *et al*., 2021; Perret *et al*., 1990); loss of m^1^G37 results in uncharged tRNA that stalls the ribosome at m^1^G37-dependent codons (Masuda *et al*., 2021), potentially also leading to cell death. In this complex biology of m^1^G37, we explored its essentiality for cell viability to elucidate a potential role in anticodon-codon pairing by removing the methylation to “expose” any deficiency that would be otherwise hidden. We performed a genome-wide suppressor screen to isolate suppressors that support cell growth when m^1^G37 is absent. This is a systematic and unbiased approach to identify the gene network that operates at the limit of cell viability. Unexpectedly, despite the association of m^1^G37 with multiple tRNAs and multiple functions, all of the suppressors in our screen are mapped to a single gene: *proS* for prolyl-tRNA synthetase (ProRS) responsible for prolyl-aminoacylation of tRNA. This exclusiveness indicates that it is the coupling of methylation with prolyl-aminoacylation that distinguishes life from death. In the *proS* suppressors, however, this coupling is disrupted, enabling us to examine anticodon-codon pairings by prolyl isoacceptors that are now “charged” but “not methylated”. We show that a minor prolyl isoacceptor, which is normally dispensable when m^1^G37 is present, has become indispensable when m^1^G37 is absent, revealing a deficiency of the anticodon-codon pairing upon removal of the methylation that needs to be addressed by the minor isoacceptor. This deficiency results from the inability of the major prolyl isoacceptor to use wobble pairings to decode a subset of codons. Thus, m^1^G37 is the determinant that resolves codon-reading bias of the major prolyl isoacceptor. We find that many bacterial pathogens of serious threat to human health do not encode the minor prolyl isoacceptor, suggesting that cell fitness and viability of these pathogens would be severely compromised in m^1^G37 deficiency.

## Results

### A genome-wide suppressor screen identifies *proS* mutations in m^1^G37 deficiency

We used *E. coli* as a genetic model, because all of the isoacceptors that are associated with m^1^G37 are well defined and experimentally tractable (Bjork and Hagervall, 2014). As with most bacterial species, *E. coli* encodes a complete set of all essential isoacceptors, as well as an additional set of non-essential tRNAs, which together assemble into a more-than-sufficient ribosome-tRNA apparatus that readily translates all of the sense codons of the genetic code. In bacteria, m^1^G37 is synthesized by the conserved TrmD methyl transferase that is also essential for cell viability (Bystrom and Bjork, 1982) (Figure 1A). This essentiality suggests that, to generate an m^1^G37-deficient condition, a simple *trmD*-KO cannot be made. We therefore created a modified *E. coli trmD-*KO strain, in which the chromosomal *trmD* was replaced by the antibiotic Kan marker, while cell survival was maintained by expression of the gene under the control of an arabinose (Ara)-inducible promoter from a plasmid that had a temperature-sensitive (*ts*) origin of replication (Figure 1B). This strain was grown overnight in the presence of Ara at the permissive temperature 30 °C, and was then diluted into an Ara-free medium at the non-permissive temperature 43 °C for multiple cycles of growth and dilution. This cycling between growth and dilution at 43 °C was designed to irradicate the *trmD*-carrying plasmid and to eliminate residual levels of the TrmD enzyme and its m^1^G37-tRNA products. After three cycles of growth and dilution (∼20 h) at 43 °C, when cell counts were no longer visible indicating cell death, an aliquot of the culture was streaked on an Ara-free plate and incubated at 43 °C to screen for suppressors. Importantly, to minimize a potential bias of the screen, due to differences in the genetic composition of the *E. coli* host, we performed the screen in parallel using three well-characterized strains (see Methods), each constructed and grown in the same modified *trmD-*KO condition. We isolated 5 colonies from MG1655, 4 colonies from BW25113, and 3 colonies from XAC. Each isolate was streaked-purified, confirmed for the loss of the maintenance plasmid (Figure S2A), and subjected to whole-genome sequencing analysis.

**Figure 1.**
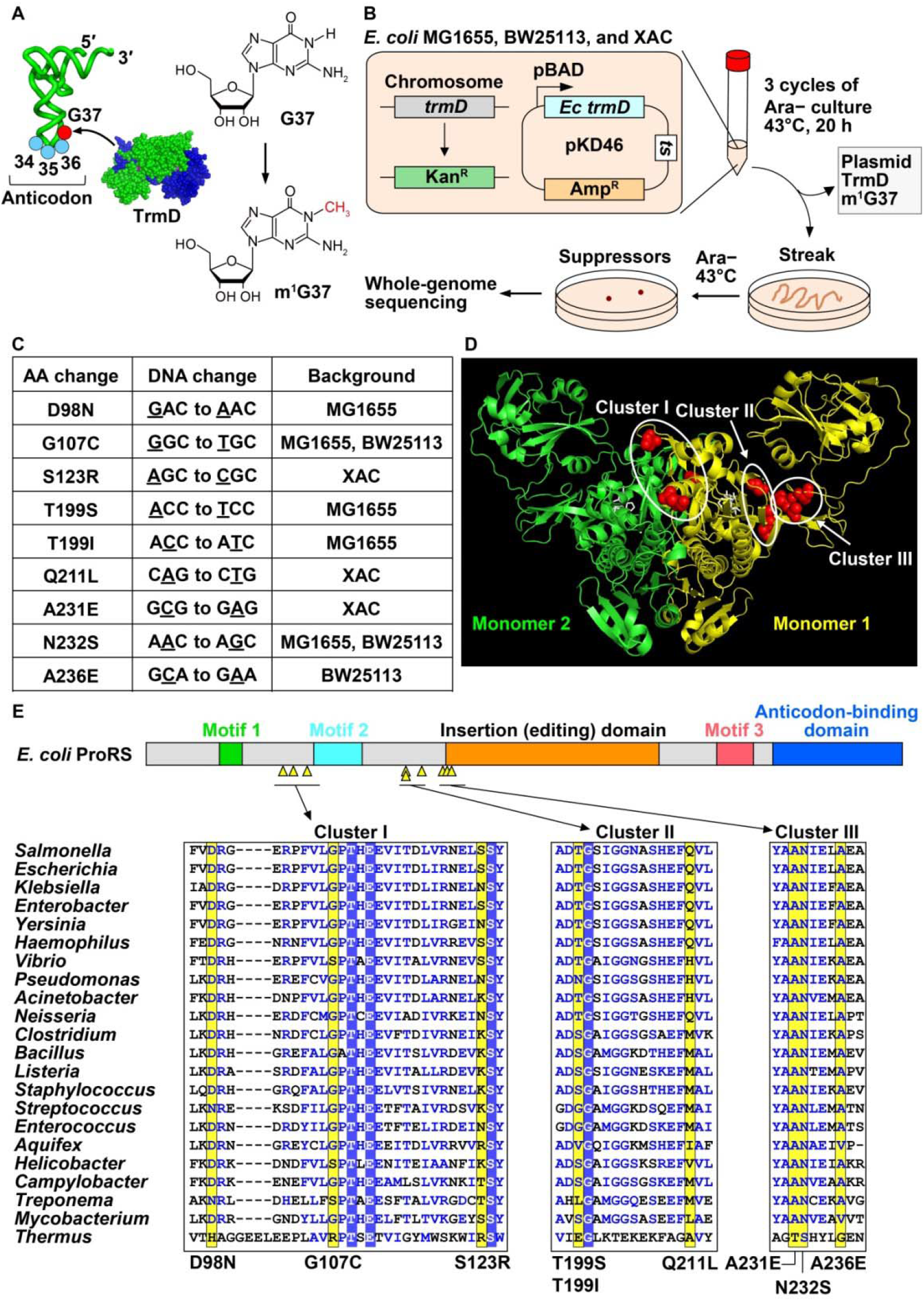
Whole-genome sequencing analysis revealed *proS* suppressor mutations of the *E. coli trmD*-KO strains. **(A)** Synthesis of m^1^G37 (red circle) by the bacterial TrmD enzyme on the 3’-side of the anticodon located at positions 34-36 (blue circles) of tRNA. **(B)** Three *E. coli trmD*-KO strains (MG1655, BW25113, and XAC) for isolation of suppressors of m^1^G37 deficiency. In each strain, the chromosomal gene was replaced with a Kan marker by P1 transduction and cell viability was supported by a plasmid-borne *trmD* under the control of the Ara-inducible pBAD promoter and a *temperature-sensitive* (*ts*) origin of replication. Culturing each strain in an Ara-free LB medium at 43 °C cured the plasmid and depleted the *trmD* enzyme and m^1^G37-tRNA. After three cycles of growth and dilution at 43 °C, cells were streaked on an Ara-free LB plate and incubated at 43 °C until appearance of suppressor colonies, each of which was streak-purified, verified for the absence of the *ts* plasmid, and subjected to whole-genome sequencing. **(C)** Exclusive mapping of suppressor mutations to *proS*, totaling in 9 unique mutations. Each unique mutation is listed for the *E. coli* strain(s) from which it was isolated, the nucleotide substitution in the chromosome, and the corresponding change of the amino acid. **(D)** Three clusters of suppressor mutations superimposed on the crystal structure of the *Enterococcus faecalis proS* enzyme in complex with the prolyl adenylate analog 5’-O-(N-(L-prolyl)-sulfamoyl) adenosine (PDB: 2J3L). In the obligate dimer structure of the enzyme, mutations are mapped to residues shown in red spheres in monomer #1, while the prolyl analog is shown in white. **(E)** A linear diagram of the structural domains of the *E. coli proS* enzyme (top) is shown with the 9 unique suppressor mutations by yellow triangles. A multi-sequence alignment of the enzyme among bacterial species (bottom) displays residues harboring mutations in yellow. Amino acids boxed in blue are conserved, while those in blue letters share structural similarity.

To our surprise, all but one clones mapped their single-nucleotide substitutions to the gene *proS*, totaling in 9 unique mutations (Figure 1C). The exception was a single clone from BW25113 that did not yield a clear whole-genome sequencing result. The exclusive mapping of all suppressor mutations to *proS*, occurring across the three strains, indicates a connection between m^1^G37 methylation and prolyl-aminoacylation of tRNA that is required for cell viability. In this connection, m^1^G37 promotes efficient prolyl-aminoacylation to generate Pro-charged tRNA^Pro^ isoacceptors that support ribosome translation of Pro codons. This connection is consistent with our published work showing that loss of m^1^G37 reduces prolyl-aminoacylation, causing ribosome stalling at Pro codons and arresting cell growth (Masuda *et al*., 2021). Thus, while m^1^G37 is associated with multiple tRNAs in multiple aspects of translation, it is most intimately associated with prolyl-aminoacylation of tRNA at the nexus that determines cell death and survival.

Mapping of each unique suppressor mutation to a known crystal structure of the *proS* enzyme in complex with a prolyl analog identified three clusters (Yaremchuk et al., 2000) (Figure 1D, E). Cluster I mutations (D98N, G107C, S123R) are localized between motif 1 and motif 2 of the catalytic domain, which is formed at the monomer-monomer interface of the obligated dimer structure of the enzyme (Yaremchuk et al., 2001) (Figure S2B). Cluster II mutations (T199S, T199I, and Q211L) are between the catalytic motif 2 and an insertion domain (also known as the editing domain) (Beuning and Musier-Forsyth, 2000; Vargas-Rodriguez and Musier-Forsyth, 2013), the latter of which provides a proof-reading activity to remove mis-acylated tRNA^Pro^. Cluster III mutations (A231E, N232S, and A236E) are near the start of the insertion domain. Some mutations in clusters II and III are also adjacent to the Pro-binding pocket of the enzyme (Figure S2B). Thus, all mutations are mapped to functionally important regions of *proS*. Additionally, all mutations are mapped to highly conserved positions of *proS* enzymes across a wide range of bacterial species (Figure 1D), indicating that they arose from the necessity to restore cell viability, even against a strong selective pressure of sequence conservation.

Each unique *proS* suppressor mutation was reconstructed by genome editing in the native MG1655 strain (Figure 2A), expressing *trmD*. In each edited strain, the chromosomal *trmD* was then deleted and confirmed for the loss of the *trmD* enzyme (Figure S2C). Note that the symbols (+) and (−) indicate the presence and absence of *trmD,* respectively. Importantly, all reconstructed suppressors supported cell viability (Figure 2B), validating that each is a bona fide suppressor of the loss of viability upon *trmD*-KO. As shown for the *proS*-D98N and *proS*-A231E suppressors (Figure 2C, Figure S2D), however, each had reduced fitness over time in a growth competition assay with the WT strain. This reduced fitness was correlated with an increased frequency of ribosomal +1 frameshifting in a reporter assay based on expression of the nLuc gene that carried a frameshift-prone insertion (Figure 2D). Thus, consistent with the notion that m^1^G37 ensures accuracy of the translational reading frame (Gamper *et al*., 2021a; Gamper *et al*., 2021b; Gamper *et al*., 2015), these m^1^G37-lacking suppressors are prone to +1 frameshifting, providing an explanation for their compromised cell fitness relative to the *trmD*-expressing WT strain. The slow fitness loss of these suppressors over the course of 24 h (Figure 2C) is likely due to the generally low efficiency of +1 frameshifting in the absence of m^1^G37 (Gamper *et al*., 2015).

**Figure 2.**
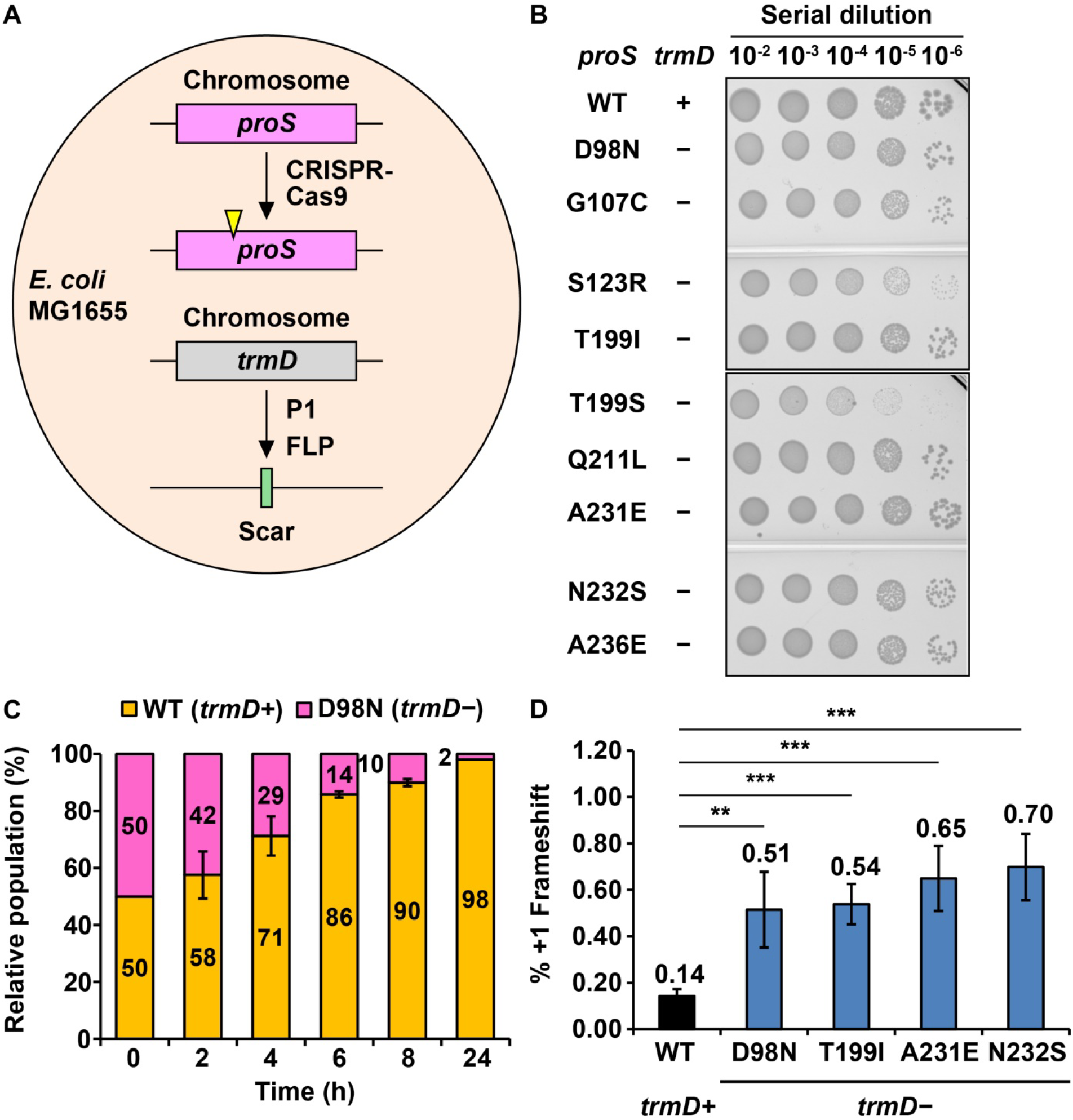
Each *proS* suppressor mutant supports cell viability without *trmD*. **(A)** Reconstruction of each *proS* suppressor mutant in MG1655. Each mutation was introduced to the chromosome by CRISPR-Cas9, followed by replacement of *trmD* with a Kan marker using P1 transduction, and by removal of the Kan marker using FLP recombination. **(B)** Viability assay of each reconstructed *proS* suppressor mutant upon a serial dilution on an LB plate incubated at 37 °C. WT is represented by the native MG1655 strain, while each suppressor is indicated by the mutation in the *proS* gene. The symbols (+) and (−) indicate the presence and absence of the *trmD* gene on the chromosome. **(C)** A cell fitness assay of a 1:1 mixture of the MG1655 WT strain (labeled with an mCherry-expressing plasmid) and the *proS*-D98N suppressor strain (labeled with a YFP-expressing plasmid) monitored over time at 37 °C in LB. Relative fraction of each strain at a given time point is based on the number of colonies of each as visualized by the fluorescence color of mCherry or YFP. **(D)** A +1-frameshifting assay measuring expression of a plasmid-borne nLuc reporter in the MG1655 WT strain or a reconstructed *proS* suppressor strain. Each strain expresses the reporter harboring an in-frame CCC triplet or a +1-frame CCC-C quadruplet next to the initiation codon AUG. The frequency of +1 frameshifting is the ratio of the reporter expression harboring the CCC-C quadruplet relative to the CCC triplet, each normalized by an internal expression of mCherry. Data are represented as mean ± SD (*n* = 4). **, *p* < 0.05; ***, *p* < 0.01.

### Decoupling of m^1^G37 methylation from tRNA aminoacylation in *proS* suppressors

As we showed recently (Masuda *et al*., 2021), the WT *proS* enzyme requires the presence of m^1^G37 for efficient prolyl-aminoacylation of tRNA^Pro^, indicating a mechanistic coupling between tRNA methylation and prolyl-aminoacylation. The isolation of *proS* suppressors that are viable but lack m^1^G37 suggests that this coupling is disrupted, such that each suppressor restores prolyl-aminoacylation without the need for m^1^G37. We tested this possibility by probing the intracellular prolyl-aminoacylation status of each *proS* suppressor, lacking *trmD* (Figure 3A). Total tRNA was isolated from each suppressor and the prolyl-aminoacylation status at the time of cell harvest was monitored for the Pro(UGG) isoacceptor using a tRNA-specific oligonucleotide. This experiment was performed in an acid condition to preserve the charged state, starting with isolation of total tRNA to separation of the charged from the uncharged states on a denaturing gel (Masuda *et al*., 2021). The prolyl-aminoacylation status was calculated by the band intensity of the charged state over the sum of both the charged and uncharged state. The result showed that the prolyl-aminoacylation status was generally high across all of the *proS* suppressors, ranging from 63% for the A231E (−) suppressor to 38% and 42% for the T199S (−) and S123R (−) suppressors (Figure 3A). The high charging status of the A231E (−) suppressor at 63% is comparable to the status of 73% of the WT (+) strain that expresses *trmD*.

**Figure 3.**
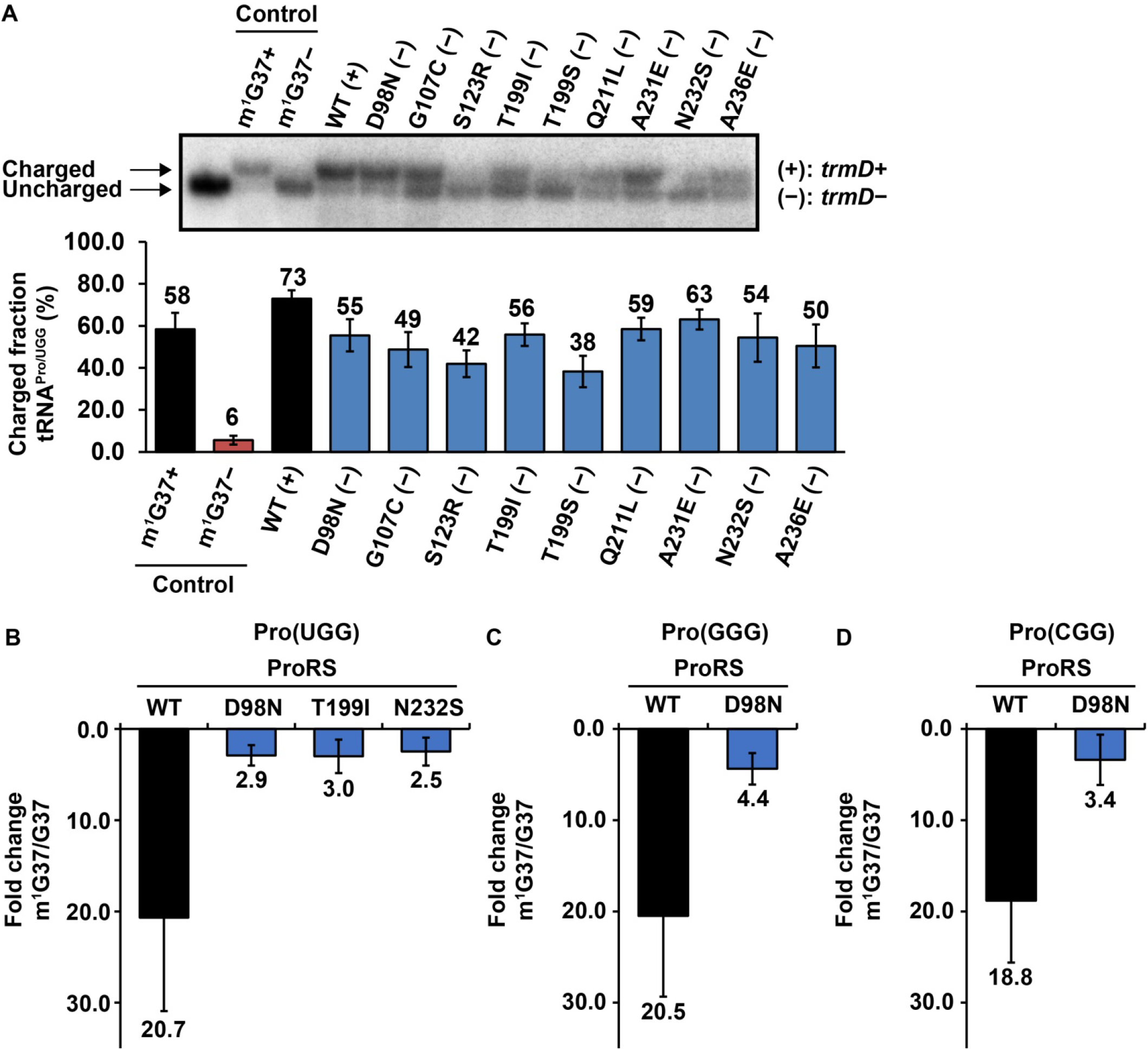
*proS* suppressor mutants are less dependent on m^1^G37 for prolyl-aminoacylation of tRNA. **(A)** Acid urea gel analysis of the prolyl-aminoacylation status of each reconstructed *proS* suppressor strain. Total RNA extracted in an acidic condition from each strain in early-log phase was run on a denaturing 12% PAGE/7M urea gel at pH 5.0 and probed for the Pro(UGG) tRNA by Northern blots. A pair of control tRNA samples isolated from m^1^G37(+) and m^1^G37 (−) conditions yielded a prolyl-aminoacylation status of 58% and 6%, respectively, whereas all reconstructed *proS* suppressors showed a status significantly higher than the negative control, ranging in 38-63%, indicating the ability to perform prolyl-aminoacylation in the absence of m^1^G37. **(B)** Kinetics of prolyl-aminoacylation on the isoacceptor Pro(UGG), comparing the loss of *k*_cat_/*K*_m_ of the WT enzyme upon removal of m^1^G37 (from the m^1^G37-state to the G37-state) relative to three of the reconstructed suppressor mutants *proS*-D98N, T199I, and N232S. **(C)** Kinetics of prolyl-aminoacylation on the isoacceptor Pro(GGG), comparing the loss of *k*_cat_/*K*_m_ of the WT enzyme relative to the suppressor mutant *proS*-D98N. **(D)** Kinetics of prolyl-aminoacylation on the isoacceptor Pro(CGG), comparing the loss of *k*_cat_/*K*_m_ of the WT enzyme relative to the suppressor mutant *proS*-D98N. All data are represented as mean ± SD. (*n* = 3). See Figure S3 for detailed kinetic parameters.

Notably, both a negative and a positive control of total tRNA were isolated from a separately constructed *trmD*-KD strain (Gamper *et al*., 2015), where the chromosomal *trmD* was removed, while a plasmid-expressed human counterpart gene *trm5* under the control of an Ara-inducible promoter was used to maintain cell viability. In this *trmD*-KD strain, addition of Ara to the medium induced the m^1^G37+ condition, producing a charging status of Pro at 58%, while depletion of Ara from the medium induced the m^1^G37− condition, producing a charging status at 6% (Figure 3A). The choice of human *trm5* as the maintenance gene was because the protein product is rapidly degraded upon turning off the Ara-controlled promoter (Christian et al., 2013), thus immediately generating the m^1^G37− condition. In contrast, a *trmD*-KD strain maintained by *E. coli trmD* would take much longer time to generate an m^1^G37− condition (Masuda *et al*., 2021), due to the high stability of the protein product in cells. We found that, relative to these controls, even the low charging status of the T199S (−) and S123R (−) suppressors (at 38-42%) is considerably higher than the charging status of the negative control and is within 2-fold of the positive control (Figure 3A). Thus, all of the isolated *proS* suppressors maintain a high intracellular prolyl-aminoacylation status, despite the absence of m^1^G37.

We next probed the enzymatic prolyl-aminoacylation activity of each *proS* suppressor mutant, using *in vitro* kinetic assays to generate intrinsic parameters that define the catalytic efficiency of each enzyme. These kinetic assays served as an independent approach from cell-based assays to verify the decoupling of aminoacylation from m^1^G37 methylation. They also offered the kinetic sensitivity to distinguish among all prolyl isoacceptors for the level of decoupling. The *E. coli* genome expresses a complete set of prolyl isoacceptors, consisting of Pro(UGG), Pro(GGG), and Pro(CGG). To focus on the effect of m^1^G37, each isoacceptor was prepared in the G37-state and in the m^1^G37-state, differing in the presence and absence of the methylation. The G37-state was transcribed from each gene with natural nucleotides, lacking m^1^G37 and any other modification, whereas the m^1^G37-state was transcribed and was post-transcriptionally modified by TrmD with the single methylation (Gamper *et al*., 2015).

Representative members of *proS* suppressor enzymes, each purified in the recombinant form to near homogeneity, were assayed for prolyl-aminoacylation with all three isoacceptors. The assay was performed in multiple-turnover condition as a function of tRNA concentration, yielding kinetic parameters *K*_m_ (the concentration of the tRNA that produces half-maximum of the enzyme activity), *k*_cat_ (the catalytic turnover), and *k*_cat_/*K*_m_ (the catalytic efficiency per tRNA-binding event). For prolyl-aminoacylation of the Pro(UGG) isoacceptor, while the *k*_cat_/*K*_m_ of the WT enzyme decreased by 20.7-fold upon loss of m^1^G37, it decreased by no more than 3-fold for three *proS* suppressor enzymes (Figure 3B, Figure S3A). Specifically, the decrease was by 2.9-fold for the *proS*-D98N enzyme (a member of cluster I), by 3.0-fold for the *proS*-T199I enzyme (a member of cluster II), and by 2.5-fold for the *proS*-N232S enzyme (a member of cluster III). The large decrease of *k*_cat_/*K*_m_ of the WT enzyme upon loss of m^1^G37 is consistent with the decrease we observed previously (Masuda *et al*., 2021), confirming the dependence on the methylation for aminoacylation, whereas the much smaller decrease of *k*_cat_/*K*_m_ of the three suppressor enzymes indicates a loss of the dependence. Thus, across all three clusters of *proS* suppressor mutants, prolyl-aminoacylation is no longer tightly coupled to m^1^G37 methylation.

The loss of coupling was also observed for prolyl-aminoacylation of the other two isoacceptors, using the *proS*-D98N mutant enzyme as an example. For prolyl-aminoacylation of the Pro(GGG) isoacceptor, loss of m^1^G37 decreased *k*_cat_/*K*_m_ by 4.4-fold for the mutant enzyme, but by 20.8-fold for the WT enzyme (Figure 3C, Figure S3B). Similarly, for prolyl-aminoacylation of the Pro(CGG) isoacceptor, loss of m^1^G37 decreased *k*_cat_/*K*_m_ by 3.4-fold for the mutant enzyme, but by 18.8-fold for the WT enzyme (Figure 3D, Figure S3C). Thus, consistent across all three isoacceptors of *E. coli*, while prolyl-aminoacylation by the WT *proS* enzyme is strongly coupled to the presence of m^1^G37, this coupling is weakened in suppressor mutants, resulting in synthesis of Pro-charged tRNA^Pro^ in the absence of m^1^G37, which underlies the basis for sustaining cell viability in *trmD*-KO.

### The Pro(GGG) isoacceptor is required for cell viability of *proS* suppressors

The prolyl-aminoacylation activity of *proS* suppressors without the need for m^1^G37 offered a way to generate Pro-charged tRNAs that were aminoacylated but not methylated. These tRNAs would then be competent to enter the ribosomal machinery, allowing us to investigate the quality of anticodon-codon pairing in the absence of the methylation. Without this prolyl-aminoacylation, none of the prolyl isoacceptors would be permitted to perform anticodon-codon pairing on the ribosome. The lack of m^1^G37 from each isoacceptor then provided a framework to identify any bias against anticodon-codon pairing. In the native state of *E. coli* with m^1^G37, each of the three prolyl isoacceptors has a dedicated responsibility for decoding. Pro(CGG) (encoded by *proK*) is dedicated for reading the CCG codon; Pro(GGG) (encoded by *proL*) is dedicated for reading the CCC and CCU (CC[C/U]) codons; and Pro(UGG) (encoded by *proM*) is capable of reading all four CCN codons, due to the presence of the cmo^5^U34 (5-caroxy-methoxyl uridine 34) modification that expands the decoding capacity to all four nucleotides (Nasvall et al., 2004; 2007) (Figure 4A, Figure S1). Of the three, the UGG isoacceptor is the major species due to its high abundance in the prolyl family (Wei et al., 2019). It is also the only one that is essential for viability in the family and cannot be deleted (Masuda *et al*., 2019; Nasvall *et al*., 2004). We therefore used the genetic tools available to *E. coli* and deleted either the GGG or the CGG isoacceptor from a *proS* suppressor strain. We then determined, upon removal of m^1^G37, if any bias against an anticodon-codon pairing could be exposed.

**Figure 4.**
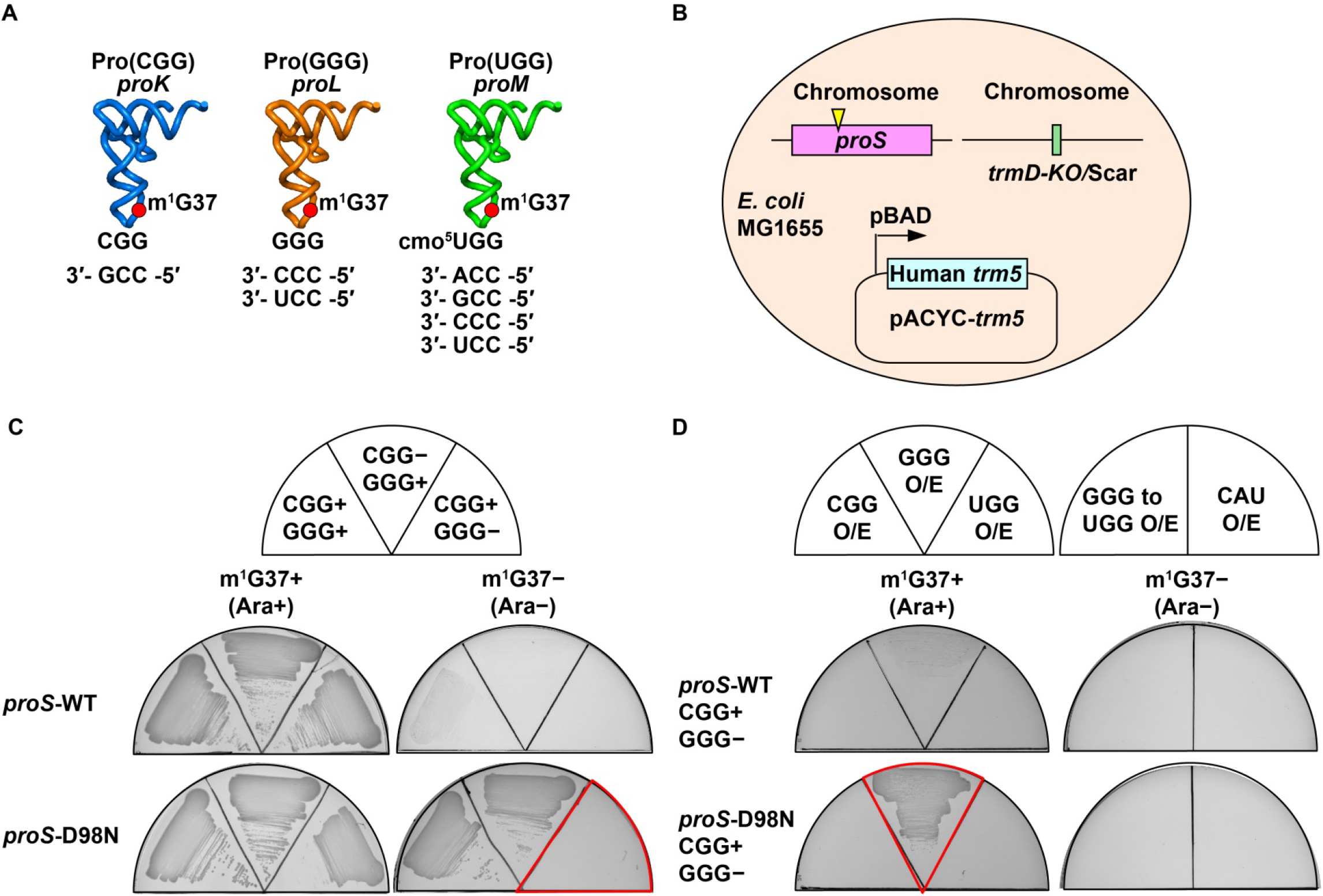
*proS* suppressor mutants require the Pro(GGG) isoacceptor for survival. **(A)** The three prolyl isoacceptors in *E. coli*, consisting of Pro(CGG) encoded by *proK* and responsible for reading the CCG codon; Pro(GGG) encoded by *proL* and responsible for reading CC[C/U] codons, and Pro(UGG) encoded by *proM* and capable of reading CCN codons due to the cmo^5^U34 modification. **(B)** Construction of a *trmD*-KD strain with a *proS* suppressor mutation. A *proS* mutation was reconstructed on the chromosome of MG1655, followed by removal of *trmD*, while cell viability was maintained by a plasmid-borne human *trm5* under the control of the Ara-inducible promoter pBAD. **(C)** Depletion analysis of Pro(GGG) or Pro(CGG) in the *trmD*-KO strain. While the strain maintained by the WT *proS* is viable in the m^1^G37+ condition, it is not viable in the m^1^G37− condition. In contrast, the strain maintained by the *proS*-D98N suppressor is viable in the m^1^G37− condition, but only when the Pro(GGG) isoacceptor is present. **(D)** Isoacceptor complementation analysis in the *trmD*-KO strain. The strain maintained by the *proS*-D98N mutation is complemented by over-expression (O/E) of the Pro(GGG) tRNA (Left), but not by over-expression of the Pro(GGG) tRNA where the anticodon is mutated to UGG, or by over-expression of the Met(CAU) tRNA. Each tRNA is expressed from the pKK223-3 plasmid.

We chose the previously constructed *trmD*-KD strain for this purpose (Gamper *et al*., 2021a; Gamper *et al*., 2015), considering that deletion of the GGG or the CGG isoacceptor in the absence of m^1^G37 might be lethal if a bias exists against an anticodon-codon pairing that would block genome-wide translation. In this *trmD*-KD strain, we reconstructed a *proS* mutation to the host genome, where *trmD* was removed from the chromosome and cell viability was maintained by expression of a plasmid-borne human *trm5* under the control of an Ara-inducible promoter (Figure 4B). The benefit of using Ara to control the biosynthesis of m^1^G37 is the ability to identify biases that could otherwise threaten cell viability in the m^1^G37− condition.

The results showed that the *trmD*-KD strain, when supported by the WT *proS*, was viable in the m^1^G37+ condition, but was non-viable in the m^1^G37− condition, consistent with the notion that m^1^G37 is required for efficient prolyl-aminoacylation to confer cell growth (Figure 4C, first row). In contrast, the *trmD*-KD strain, when supported by the *proS*-D98N suppressor, was viable in the m^1^G37− condition, but this viability was observed only when the minor isoacceptor GGG was present (second row). Thus, while the *proS*-D98N suppressor was able to support cell viability without the need for m^1^G37, it failed to do so when the GGG isoacceptor was also lacking. This dependence on the GGG isoacceptor for viability was also observed for the *proS*-T199I and *proS*-N232S suppressors (Figure S4), indicating a shared mechanism. Upon over-expression of a plasmid-borne GGG isoacceptor, however, cell viability was restored in the m^1^G37− condition (Figure 4D, second row). This restoration was not observed in the *trmD*-KD strain supported by the WT *proS* (first row), or in the *proS*-D98N suppressor supported by over-expression of a variant of the GGG isoacceptor that harbored the UGG anticodon, or in the suppressor strain supported by over-expression of a CAU isoacceptor specific for reading of the Met AUG codons (Figure 4D, right column). Thus, the *proS*-D98N suppressor requires Pro(GGG) for cell viability, implying that the lack of the isoacceptor has rendered decoding of the associated CC[C/U] codons insufficient. Normally, decoding of CC[C/U] is a shared responsibility between the minor GGG isoacceptor and the major UGG isoacceptor that is modified with cmo^5^U34 (Figure 4A). The dependence on the minor isoacceptor for viability of the *proS-*D98N suppressor implies that decoding of CC[C/U] codons is no longer supported by the major isoacceptor in the absence of m^1^G37.

### Codon usage bias for expression of cold-shock response genes

We tested the hypothesis that, in the absence of m^1^G37, translation of CC[C/U] codons would be dependent on the availability of the minor isoacceptor Pro(GGG). A genome-wide codon usage analysis of *E. coli* protein-coding genes showed an average usage of CC[C/U] at 0.31 when normalized by the sum of all Pro codons that occur in the same gene (Figure 5A). Interestingly, 20.4% of *E. coli* protein-coding genes display an average usage of CC[C/U] higher than 0.5, indicating an enrichment of these codons. In GO analysis, these CC[C/U]-enriched genes are most populated in the group involved in organization of cellular components in response to stress with a frequency of 4.2-fold above the background frequency (Figure 5B). Notably, many genes in this group encode cold-shock protein (Csp) products in response to temperature downshift (Figure S4A), possibly serving as chaperones to prevent mRNA from aggregation and to facilitate initiation of protein synthesis (Keto-Timonen et al., 2016). We tested the possibility that the high enrichment of CC[C/U] codons in *csp*-related genes would render the *E. coli* response to cold-shock dependent on Pro(GGG) when m^1^G37 is absent. Indeed, the WT strain, when supported by the native *proS* while expressing *trmD* and synthesizing m^1^G37, remained viable upon temperature shift from 37 °C to 22 °C, whereas several *proS* suppressors, lacking *trmD* and unable to synthesize m^1^G37, did not (Figure 5C). The loss of viability was notable for the *proS*-D98N mutant even with a milder temperature downshift to 30 °C (Figure S4B). The high sensitivity of these suppressors to the cold shock stress is consistent with the notion that, in the absence of m^1^G37, translation of CC[C/U]-enriched *csp*-related genes is compromised, likely due to insufficient supply of the Pro(GGG) isoacceptor.

**Figure 5.**
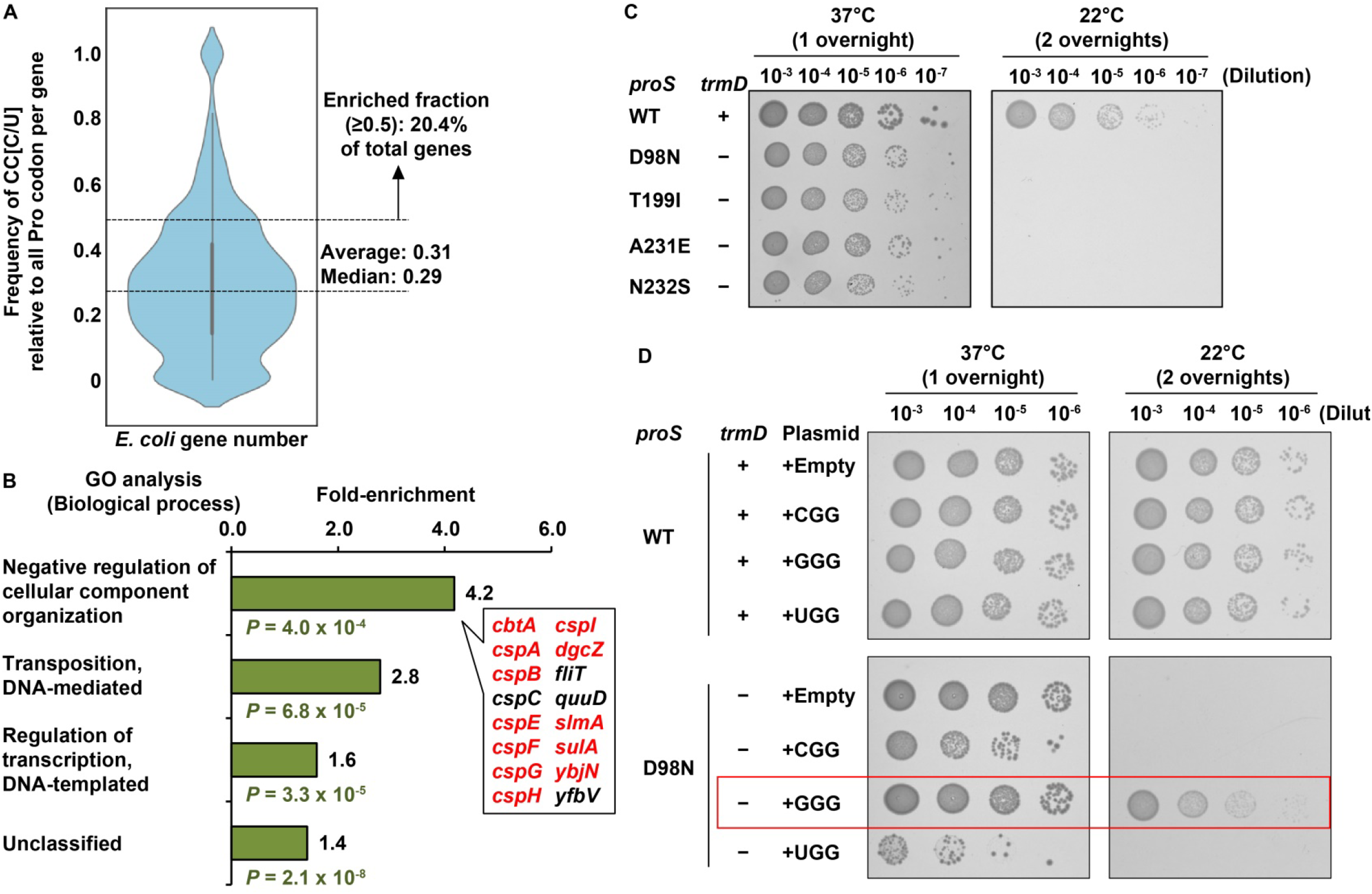
Expression of *E. coli csp* genes is sensitive to the cellular level of Pro(GGG). **(A)** A violin plot of codon usage frequency of CC[C/U] in *E. coli* protein-coding genes. About 20.4% of total protein-coding genes have a higher codon usage frequency than the average, which are used for Gene Ontology (GO) analysis. **(B)** Identification of four groups of *E. coli* protein-coding genes enriched with CC[C/U] codons. Of the 16 genes categorized in the top group, 12 (in red) are enriched with CC{C/U] codons, while 8 of the 12 are of the *csp* family. **(C)** Cold sensitivity assay. The MG1655 WT (+) strain and the reconstructed *proS*-D98N suppressor strain (−) (Figure 2A) were serially diluted and tested for viability at 37 °C (one overnight) or at 22 °C (two overnights). **(D)** Rescue of cold sensitivity by expression of tRNA isoacceptors for the reconstructed *proS*-D98N mutant. Each isoacceptor was expressed from the pKK223-3 plasmid in the WT (+) strain or the *proS*-D98N strain.

In *E. coli*, CC[C/U] codons are rare (average usage at 0.31, Figure 5A) relative to the more abundant CC[A/G] codons for Pro, raising the possibility that the intrinsic level of the corresponding ProGGG) would be similarly low relative to the others. This possibility was confirmed by over-expression of the isoacceptor from a multi-copy plasmid, showing restoration of viability at 22 °C (Figure 5D, Figure S5B). In contrast, no restoration of viability was observed by over-expression of the other two isoacceptors. The specific requirement of Pro(GGG), but not others, to restore cell viability at 22 °C in m^1^G37-lacking suppressors supports the notion that it is the GGG isoacceptor that is responsible for translation of the highly enriched CC[C/U] codons that are necessary to produce the cold-shock response.

### Absence of the Pro(GGG) isoacceptor from bacterial species

The requirement of the Pro(GGG) isoacceptor for cell viability in m^1^G37-lacking suppressors is important for fundamental understanding of the genome structure across the diversity of bacterial species. While m^1^G37 is strictly conserved throughout the bacterial domain (Bystrom and Bjork, 1982), not all bacterial species encode the minor Pro(GGG) isoacceptor. In such a simplified genome structure, decoding of CC[C/U] codons could only be achieved by the cmo^5^U34-modified major isoacceptor Pro(UGG), while decoding of the CCG codon could be achieved by both the major isoacceptor and the other minor isoacceptor Pro(CGG). We thus surveyed the genomic tRNA database (GtRNAdb) to understand expression of all prolyl isoacceptors. We compared the presence and absence of each isoacceptor with a combined 16S and 23S ribosomal RNA (rRNA) phylogeny of selected genera in GtRNAdb (Figure 6A). Nearly all examined bacterial genomes encode the major isoacceptor Pro(UGG), supporting the essentiality of the isoacceptor for cell viability, while a few exceptions lacking the isoacceptor are likely due to incomplete analysis of the genomes. Although some bacterial species also encode a minor Pro(AGG) isoacceptor, this is extremely rare in occurrence (12 occurrences across the species surveyed) and has no pattern of evolutionary conservation, suggesting that they are likely tRNA pseudogenes and are therefore not considered further. The resulting phylogenetic tree revealed a bifurcation of bacterial species into phyla with either the presence or absence of the minor Pro(GGG) isoacceptor. Notably, almost all the clades that are missing Pro(GGG) are also missing Pro(CGG), indicating a co-existence relationship between these two minor isoacceptors (Figure S5). The absence of both isoacceptors from a genome thus signifies the sole dependence on the major isoacceptor for translation of all Pro codons. This is most notable in the entire class of Mycoplasma, which are intracellular parasites with minimal sets of tRNA isoacceptors (Razin *et al*., 1998). The only exception to the co-existence rule of both minor isoacceptors is in the family Lactobacillaceae, which lack Pro(GGG) but contain Pro(CGG) (Figure 6A), an observation of interest for future studies.

**Figure 6.**
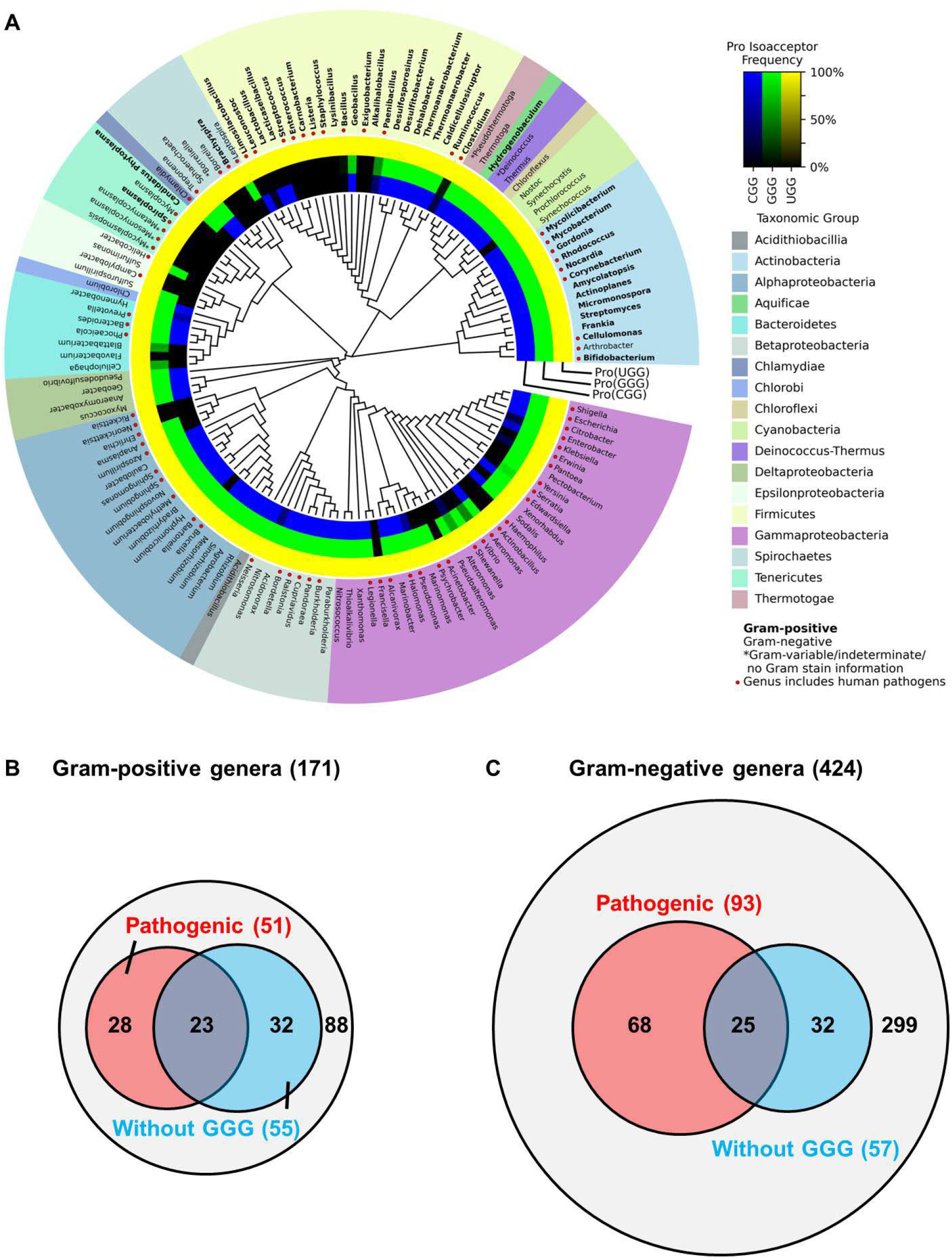
Absence of Pro(GGG) from some bacterial species. **(A)** A maximum-likelihood phylogenetic tree showing the presence and absence of prolyl tRNA genes in genomes of representative bacterial genera that are available at GtRNAdb. This is generated with the concatenated 16S and 23S alignments with higher classifications as taxonomic groups shown in various colors. Gram-positive (Gram (+)) genera are shown in bold face, and those of human pathogens are indicated by a red circle. Inner rings represent the presence and absence of prolyl isoacceptor genes: the blue ring for Pro(CGG), the green ring for Pro(GGG), and the yellow ring for Pro(UGG). The gradient of color darkness indicates the frequency of the species lacking the corresponding isoacceptor gene within the genus. Darker color means more species lacking the isoacceptor gene, with the complete black indicating the complete absence. **(B)** A Venn diagram depicting the distribution of medically relevant (pathogenic) genera with and without Pro(GGG) within Gram-positive genera. Among 171 genera, 51 genera contain pathogenic species (orange), 55 genera lack Pro(GGG (blue), and 23 are in common, indicating pathogenic species that lack Pro(GGG) (blue). **(C)** A Venn diagram depicting the distribution of medically relevant (pathogenic) genera with and without Pro(GGG) within Gram-negative genera. Among 424 genera, 93 genera contain pathogenic species (orange), 57 genera lack Pro(GGG) (blue), and 25 are in common, indicating pathogenic species that lack Pro(GGG).

Many bacterial species lacking the Pro(GGG) isoacceptor are infectious human pathogens. In Gram-positive genera, 55 of the 171 species (32%) lack Pro(GGG), and 23 of the 55 are human pathogens, including species of the phyla *Enterococcus* and *Staphylococcus* (Figure 6A, B). In Gram-negative genera, 57 of the 424 species (13%) lack Pro(GGG), and 25 of the 57 are human pathogens, including species of the phylum *Acinetobacter* (Figure 6A, C). Notably, the pathogens mentioned here constitute the ESKAPE family, posing an additional threat to human health, due to their acquisition of resistance to almost all currently available antibiotics (De Oliveira et al., 2020). The prevalence of lacking Pro(GGG) from these pathogens suggests a medical relevance that can be explored for antibiotic targeting.

## Discussion

Codon usage is well known to be closely correlated with the abundance and composition of the cellular tRNA pool (Rak *et al*., 2018). Much less known is how codon usage may be controlled or modulated by various tRNA post-transcriptional modifications. Even less known is how codon usage is determined by position-37 post-transcriptional modifications, which occur on the 3’-side of the anticodon and are not directly involved in the anticodon-codon pairing. Yet, elimination of a position-37 modification from a cell has resulted in ribosome stalling and programmatic changes of gene expression (Lamichhane et al., 2016; Masuda *et al*., 2021; Thiaville *et al*., 2016), strongly implicating that the position-37 modification modulates the quality of anticodon-codon pairing. This modulation however has never been directly tested. Here we show that the position-37 modification with m^1^G37 is an important neutralizer that addresses the deficiency in, and bias against, specific anticodon-codon pairings established by the position-34 wobble modification in the major Pro(UGG) isoacceptor. Typically, U34 of the major isoacceptor is modified into cmo^5^U34, which expands the decoding to all four nucleotides (Nasvall *et al*., 2004; 2007). We show that, while this expansion supports cell viability when m^1^G37 is present, it no longer can do so when m^1^G37 is absent (Figure 4D). Thus, m^1^G37 is the determinant that enables the cmo^5^U34-modified UGG anticodon to expand decoding, whereas lacking m^1^G37, the modified UGG anticodon is unable to read CC[C/U] codons, requiring the minor isoacceptor Pro(GGG) to assume the responsibility (Figure 7A). Notably, the minor isoacceptor Pro(GGG) is designed to only read CC[C/U] codons, although the reading is non-essential in m^1^G37 abundance, due to coverage of the codons by the cmo^5^U34-modified major isoacceptor. However, the reading of the minor isoacceptor becomes essential in m^1^G37 deficiency, emphasizing the notion that m^1^G37 is the neutralizer that resolves the deficiency of the cmo^5^U34-modified major isoacceptor to read CC[C/U] codons. Importantly, m^1^G37 is in a conserved modification circuit with the chemical moiety of cmo^5^U34 in the major isoacceptor Pro(UGG) throughout evolution. This includes the methyl addition to the cmo^5^U34 moiety in *E. coli* Pro(UGG), generating mcmo^5^U34 (5-methoxy-carbonyl-methoxy-uridine 34) in bacteria, and the ncm^5^U34 (5-carbamoyl-methyl-uridine 34) modification in eukaryotes, sharing the carbamoyl chemical moiety at the 5-position of the uridine (Figure S1). This broad conservation indicates that the functional compensation of m^1^G37 for the deficiency of cmo^5^U34-mediated decoding of Pro codons as we report here has sustained a strong selective pressure over the evolution time.

**Figure 7.**
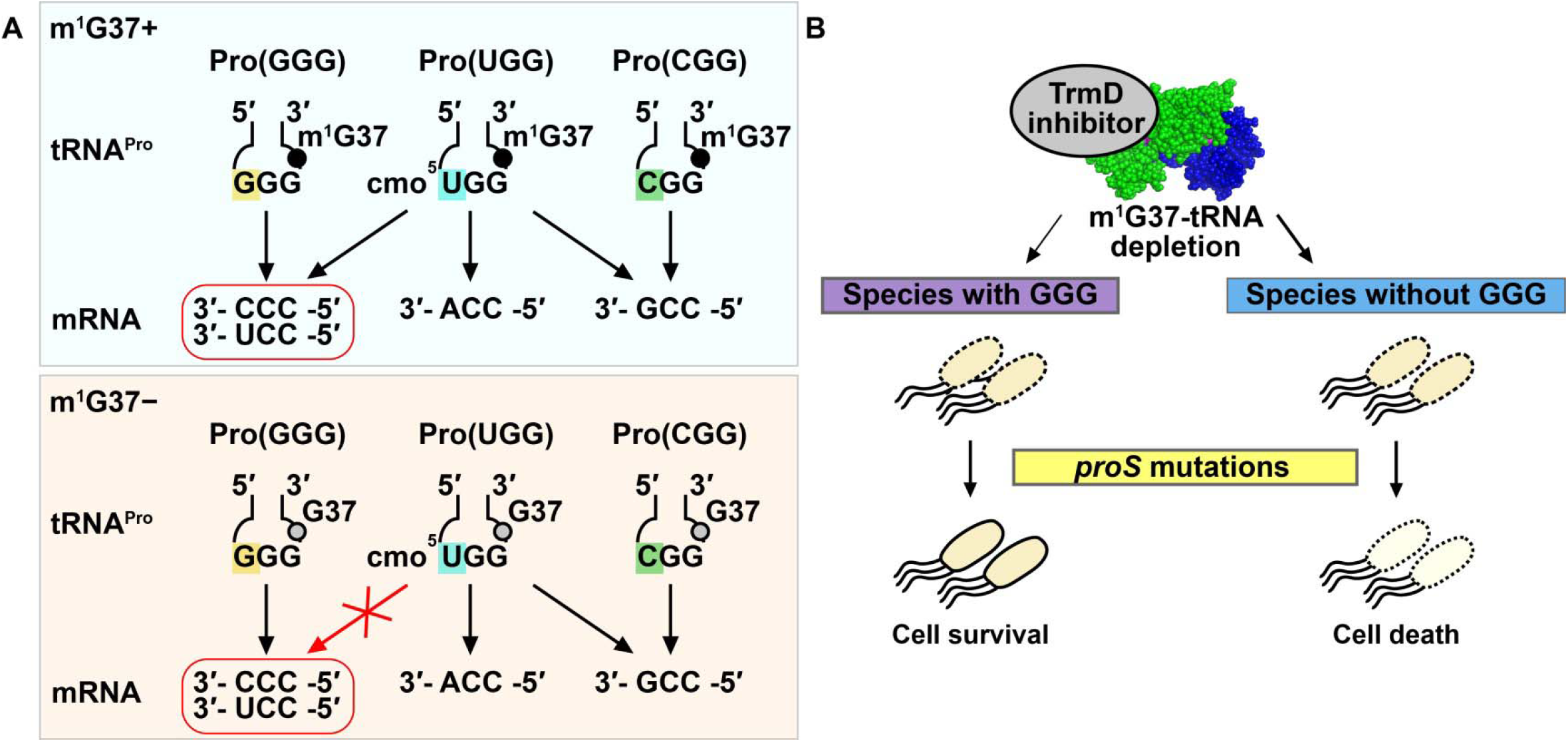
tRNA methylation with m^1^G37 resolves codon bias of the cmo^5^U34-modified major isoacceptor Pro(UGG) with medical relevance. **(A)** m^1^G37 resolves bias of the major prolyl isoacceptor. (Top) In the presence of m^1^G37 (a black circle), the major isoacceptor Pro(UGG), modified with cmo^5^U34, is capable of reading all four Pro codons, including using wobble base pairings to read CC[C/U]. (Bottom) In the absence of m^1^G37− (an open circle), the cmo^5^U34-modified major isoacceptor Pro(UGG) is no longer capable of reading CC[C/U], thus requiring the minor isoacceptor Pro(GGG) to assume the role and rendering the latter essential for cell viability. **(B)** Risk of resistance of targeting TrmD. As a high-priority antibacterial target, TrmD is being actively pursued for discovery of new antibiotics. Upon targeting TrmD by an inhibitor, while cell viability is diminished, a potential resistance mechanism could occur by suppressor mutations in *proS* as shown here. However, this resistance mechanism would be thwarted if the bacterial species lacks the Pro(GGG) isoacceptor, which becomes essential for viability in the absence of TrmD. Attractive candidates for this anti-TrmD strategy would include the Gram-negative *Acinetobacter baumannii*, and the Gram-positive *Enterococcus faecalis* and *Staphylococcus aureus*.

The ability of m^1^G37 to resolve the deficiency of the cmo^5^U34-modified major isoacceptor now provides new insight into the long-standing question of how cmo^5^U34 expands the pairing capacity to all four nucleotides. In crystal structures, the base pairing of cmo^5^U34 with C and U is mediated by just one hydrogen (H)-bond each, whereas those with A and G are mediated by three H-bonds each (Figure S1) (Weixlbaumer et al., 2007). While these structures were obtained with a synthetic anticodon stem-loop (ASL), lacking the position-37 modification, they nonetheless raise the question of how the single H-bond can provide the same stability as three H-bonds. This question has remained unsolved for decades, due to the lack of a tool to probe the question at a biological scale. Here we use a genetic tool to remove m^1^G37 and, upon removal, we find that the single H-bond in the cmo^5^U34-C/U pairing is indeed insufficient and unable to satisfy the high demand of genome-wide translation required for cell viability. In the major isoacceptor Pro(UGG), stabilization of the cmo^5^U34-C/U pairing by m^1^G37 is likely rooted in the ability of the methylation to remodel the ASL structure, which adopts a disordered form when lacking m^1^G37 (Maehigashi et al., 2014). This implication is presumably far reaching, given that the cmo^5^U34-mediated anticodon-codon pairing is wide-spread among 5 other tRNA families, in addition to Pro(UGG). In *E. coli*, cmo^5^U34 co-exists with m^1^G37 in Leu(UAG), with ms^2^i^6^A37 in Ser(UGA), with m^6^A37 in Val(UAC), and with ct^6^A37 (cyclic *N*^6^-threonyl-carbamoyl-adenosine 37) in Thr(UGU). Although cmo^5^U34 in Ala(UGC) does not co-exist with a position-37 modification, the deficiency of its pairing with C/U may be resolved by other post-transcriptional modifications in the tRNA. Further structural analysis of a cmo^5^U34-C/U base pairing in the presence of a position-37 modification, or accompanied by the full complement of all post-transcriptional modifications in a tRNA, will shed more light on how other tRNA modifications stabilize the cmo^5^U34-C/U base pairing.

While we emphasize the importance of m^1^G37 in codon-reading of prolyl isoacceptors, this does not mean that m^1^G37 is not important for other tRNAs that are associated with the methylation (e.g., isoacceptors of Leu and Arg). The distinction is that the importance of m^1^G37 in prolyl isoacceptors is established at the limit of cell viability, whereas the importance of m^1^G37 in others is not as critical. Indeed, all of the mutations in our genome-wide screen of suppressors of m^1^G37-KO are mapped to the single gene *proS*, indicating a unique dependence of m^1^G37 for prolyl-aminoacylation that is required for cell viability. In contrast, while we showed previously a similar dependence on m^1^G37 for aminoacylation of Arg(CCG) and Leu(UAG) (Masuda et al., 2021), no suppressor mutations are mapped to *argS* or *leuS*. These results suggest that, in the decision of life and death, cells choose to introduce mutations into *proS* at highly conserved and functionally important regions that would compromise activity of the enzyme in exchange for viability. A detailed analysis of selected *proS* suppressor mutants verifies that each has a kinetic deficiency relative to the WT enzyme (Figure S3), ranging in a loss of *k*_cat_/*K*_m_ from 4- to 170-fold. This kinetic deficiency is apparently small enough of a price for the trade-off to have the ability to continue synthesis of Pro-charged tRNA^Pro^ to support protein synthesis and cell viability. Nonetheless, the trade-off does incur a fitness cost. One reason is the increased propensity of ribosomal +1 frameshifting (Figure 2D), consistent with the notion that loss of m^1^G37 induces the ribosome to shift to the +1-frame, resulting in changes of the reading frame and cell death (Gamper *et al*., 2021a; Gamper *et al*., 2015).

The dependence of *proS* suppressors on the minor isoacceptor Pro(GGG) for cell viability has important medical implication (Figure 7B). The methyl transferase TrmD has long been ranked as a high-priority antibacterial target (White and Kell, 2004), due to its essentiality for bacterial viability, conservation across the bacterial domain, fundamental distinction from its human counterpart Trm5 in structure and mechanism (Ahn et al., 2003; Christian et al., 2004; Christian and Hou, 2007; Christian et al., 2010; Christian et al., 2016; Goto-Ito et al., 2008; Goto-Ito et al., 2009; Sakaguchi et al., 2014), and possession of a small-molecule binding pocket for the methyl donor that can be targeted by new compounds. Notably, the methyl-donor binding pocket in TrmD is unique, adopting a rare protein topological knot-fold that is absent from the great majority of methyl transferases (Ahn *et al*., 2003; Christian *et al*., 2016; Elkins et al., 2003), indicating the potential of bacteria-specific drug targeting in novel chemical space and diversity. The importance of targeting TrmD is further emphasized by our finding that m^1^G37-tRNA is required for translation of membrane-associated genes in the bacterial cell-envelop structure (Masuda *et al*., 2019), due to an enrichment of CC[C/U] codons in these genes, suggesting that targeting can disrupt the membrane-controlled mechanisms to facilitate drug entry and intracellular retention for faster bactericidal action. Indeed, developing TrmD inhibitors is currently an active pursuit (Hill et al., 2013; Thomas et al., 2020; Whitehouse et al., 2019; Zhong et al., 2019a; Zhong et al., 2019b). However, in any antibacterial strategy that targets a single gene, the risk of resistance is a major concern. The discovery of *proS* suppressors in *trmD-*KO raises the concern of a potential resistance mechanism. This concern is now ameliorated by our subsequent discovery that these suppressors require the minor isoacceptor Pro(GGG) for survival. The prevalence of lacking the minor isoacceptor in many clinically relevant bacterial pathogens (Figure 6A) suggests that these pathogens would be attractive candidates for targeting *trmD* in a new antimicrobial strategy that would not encounter a resistance mechanism due to suppressor mutations of *proS*.

A recent study reported the isolation of suppressors of m^1^G37 deficiency from several *E. coli* strains (Clifton et al., 2021), each harboring a chromosomal mutation in *trmD*. Although the majority of the isolated suppressor mutations are also mapped to *proS*, with some overlapping with the suppressors we report here, the approach and focus of the recent study is fundamentally different from ours (Table S2). Specifically, the recent study was set out to isolate suppressors of *trmD*-KD and concluded that mutations of *proS* constitute the dominant mechanism that repairs the deficiency of m^1^G37. In contrast, we use *trmD*-KO to eliminate m^1^G37, so as to isolate suppressors that would expose differential codon-reading by various tRNA isoacceptors that would have been otherwise hidden by the presence of m^1^G37. From the exposed differential codon-reading upon deletion of m^1^G37, we gain insight into how the methylation resolves the bias. We show that, while m^1^G37 is normally associated with decoding of Pro, Leu, and Arg codons in the genetic code, it is most critically associated with resolving codon bias against decoding of pyrimidine-ending codons of Pro, serving as an essential neutralizer that operates at the limit of cell survival. This discovery has important medical implication for antibacterial targeting of bacterial pathogens that are enriched with pyrimidine-ending codons of Pro and are also lacking the minor isoacceptor of Pro(GGG).

## STAR METHODS

Detailed methods are provided in the online version of this paper and include the following:

## SUPPLEMENTAL INFORMATION

Supplemental Information includes five figures and can be found with this article online.

## ACKNOWLEDGEMENTS

We thank Sean Moore for advice on isolation of suppressors, Roy Kishony for plasmid pZS2R, Glenn Bjork for antibodies to *E. coli* TrmD, and Howard Gamper for transcripts of isoacceptors of *E. coli* tRNA^Pro^. This work is supported by grant award R35 GM134931 to YMH and R01 HG006753 to TML from the National Institutes of Health. We thank Cancer Genomics and BioComputing of Complex Diseases lab at Bar-Ilan University for valuable discussions.

## AUTHOR CONTRIBUTIONS

I.M. constructed the *trmD*-KO strain and performed the genome-wide suppressor screen, Y.Y. and T.C. performed kinetic assays of prolyl-aminoacylation, R.D., S.T., R.M., and M.F.M. performed whole-genome sequencing analysis and variant calling, H.M. and T.L. performed phylogenetic analysis, S.M. generated data of acid-urea gel analysis, Y.N. performed codon usage analysis. I.M. prepared figures, legends, and methods, and Y.M.H. wrote the manuscript. All authors have given approval to the final version of the manuscript.

## DECLARATION OF INTERESTS

The authors declare no competing financial interests.

## Supplemental Information

### Key Resources Table

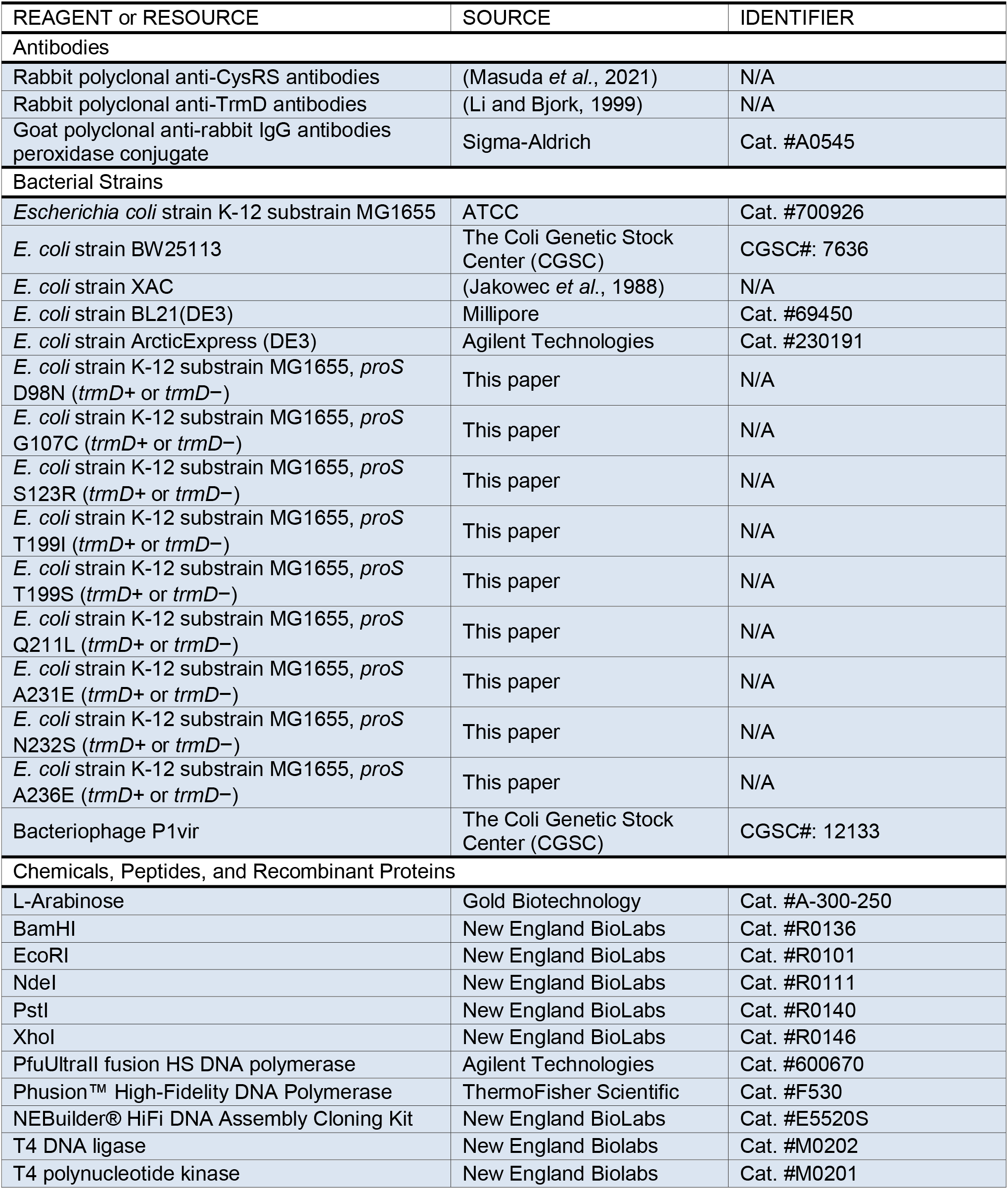
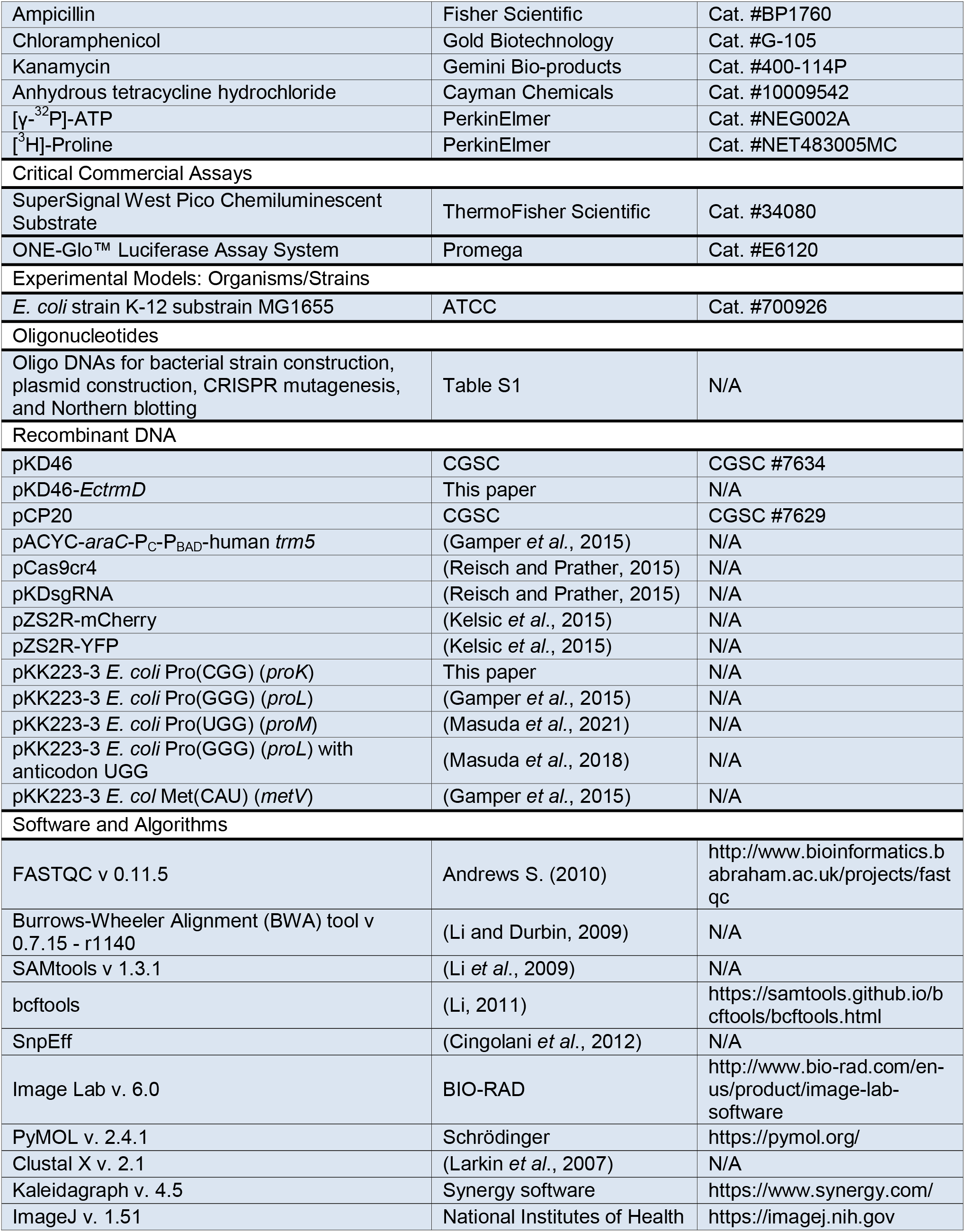
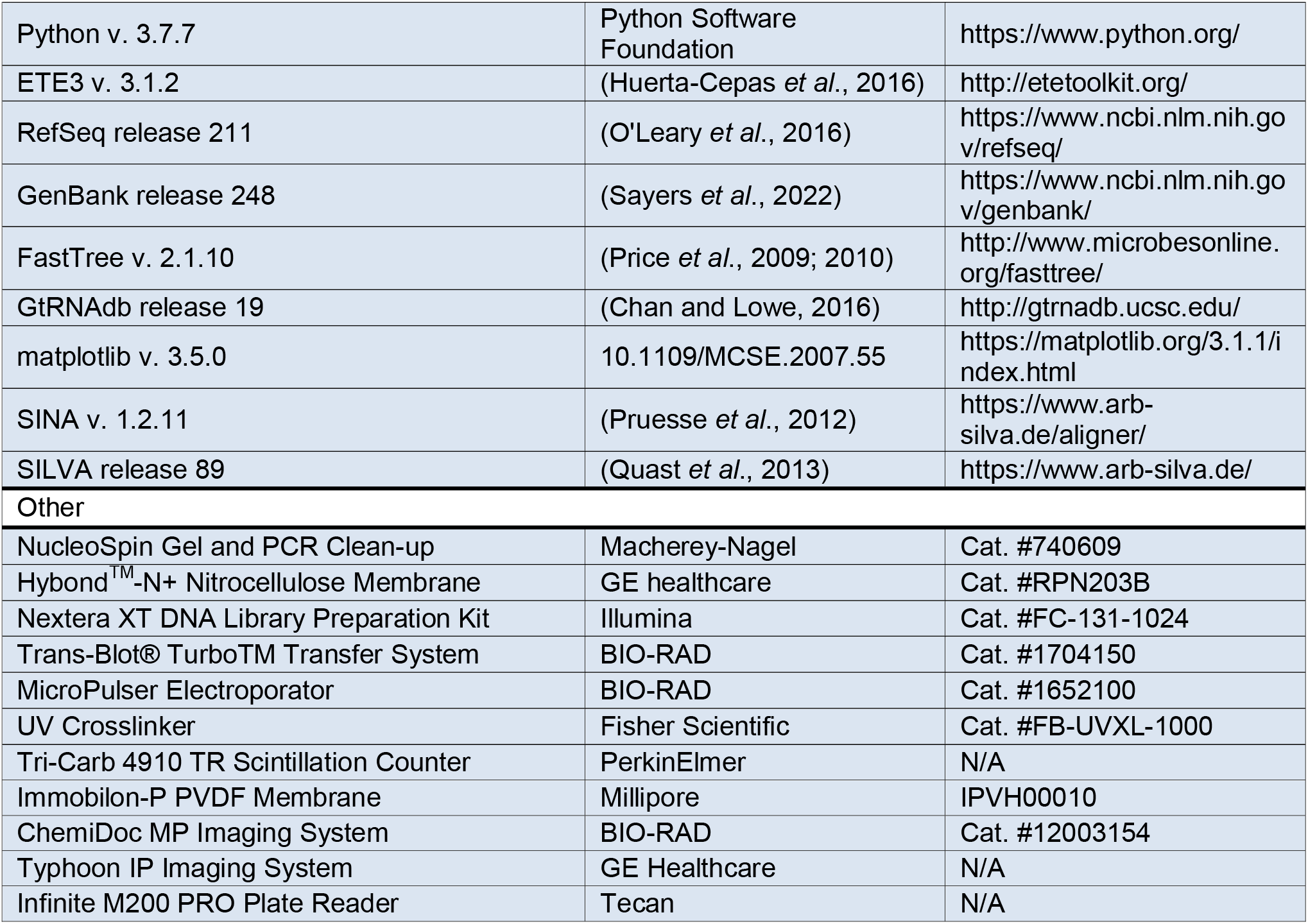

### Method Details

#### Genome-wide screen of *trmD-KO* suppressors

Because *trmD* is essential for viability, a simple KO cannot be made. A maintenance plasmid carrying *E. coli trmD* was made with a *ts* origin of replication. The gene together with a partial sequence of *P*_BAD_ was isolated from pACYC-*araC*-P_C_-P_BAD_-*EctrmD* (Gamper et al., 2015; Masuda *et al*., 2019) by excision with BamHI and XhoI (NEB) and gel purified by NucleoSpin Gel and PCR Clean-up (Macherey-Nagel). The *ts* origin of replication of pKD46 (Datsenko and Wanner, 2000) was amplified with PfuUltraII fusion HS DNA polymerase (Agilent Technologies) between the BamHI and an XhoI sites, excluding the λRed recombinase coding sequence. The PCR product was digested by BamHI and XhoI, then gel-purified by NucleoSpin Gel and PCR Clean-up (Macherey-Nagel). The *trmD* gene and the amplified region of pKD46 were ligated by T4 DNA ligase (NEB) and transformed into competent cells of one of the three *E. coli* strains of this work. The ligated plasmid (pKD46-*EctrmD*) was verified by Sanger sequencing.

The three *E. coli* strains in this work were MG1655 (Hayashi *et al*., 2006), BW25113 (Grenier *et al*., 2014), and XAC (Dincbas *et al*., 1999). After transformation with pKD46-*EctrmD,* each strain was made into *trmD*-KO by P1 transduction to replace *trmD* with a Kan marker using the P1 lysate of the *trmD*-KO strain (Gamper *et al*., 2015; Masuda *et al*., 2019). Each strain was grown overnight at 30 °C in LB with the presence of Ara 0.2% (w/v), ampicillin (Amp) (100 µg/mL), and Kan (50 µg/mL), allowing stable replication and expression of pKD46-*EctrmD* from the pBAD promoter. Cells were diluted at 1:100 to fresh LB + Kan and grown at 43 °C until OD_600_ = 1.0 for the first cycle, during which expression of *trmD* was turned off, replication of the *ts*-plasmid was inhibited, and cellular TrmD and m^1^G37-tRNA was reduced with each cell division. Cell cultures at OD_600_ = 1.0 were diluted (1:20) and passed through two more rounds of growth and dilution to pre-warmed LB + Kan (at 1:20 each) until cell counts were not visible (∼4 h). An aliquot (10 µL) of each culture was spread on an LB + Kan plate and incubated at 43 °C until distinct colonies were formed. In total, 5 colonies of MG1655, 4 colonies of BW25113, and 3 colonies of XAC were isolated, streak-purified at 43 °C, verified for the absence of pKD46-*EctrmD*, and subjected to whole-genome sequencing.

#### Whole-genome sequencing and identification of variants

For whole-genome sequencing analysis, we chose two suppressor colonies from each strain. After streak-purification, each was grown in 3 mL LB + Kan at 37 °C overnight and cells were harvested and resuspended in 500 µL of 50 mM NaCl and 25 mM EDTA. Cell lysis was performed with the addition of lysozyme (1 mg/mL) at 37 °C for 30 min, and further by the addition of 1.5% sarcosyl for 10 min at 37 °C. Cell lysates were extracted by phenol twice, phenol-chloroform-isoamyl alcohol once, and ethanol precipitated. Chromosomal DNA was re-suspended in 200 µL TE buffer (pH = 8.0) and was used to generate a genomic library using Nextera XT DNA Library Prep Kit (Illumina). A DNA library from each WT was also prepared as a reference. Each library was paired-end sequenced by MiSeq at 2 x 150 bp, yielding 80x coverage of reads relative to the entire genome.

Sequencing reads were pre-processed using FASTQC (Version 0.11.5) (Andrews, 2010), and mapped to the *E. coli* BW25113 genome sequence (Grenier *et al*., 2014) using BWA (Version 0.7.15 - r1140) (Li and Durbin, 2009) and SAMtools (Version 1.3.1) (Li *et al*., 2009). Single nucleotide polymorphisms (SNPs) in each sample were retrieved as Variant Call Format (VCF) files. A comparative analysis of SNPs between all samples with the BW25113 genome was then performed using bcftools (Li, 2011). Further annotation of SNPs for each comparison was performed using SnpEff (Cingolani *et al*., 2012). Finally, SNPs with missense and frameshift variants were considered to generate final Excel files. The SNPs only present in the suppressor genome but not in the parental WT genome were identified as variants.

Of the 6 suppressor colonies subjected to whole-genome sequencing, 5 contained a single nucleotide substitution that was mapped to *proS*, while one was unclear in sequence analysis. We therefore focused on the *proS* gene, PCR-amplified it from all suppressor clones, and performed Sanger sequence analysis of the amplified gene. We confirmed that all suppressor clones contained a single nucleotide substitution in *proS*, noting that G107C and N232S were identified independently from both MG1655 and BW25113.

#### Structural mapping and amino-acid sequence alignment

Each suppressor mutation was mapped to the crystal structure of *E. faecalis* ProRS (PDB 2J3L) (Crepin *et al*., 2006) using PyMOL (Schrödinger). The amino acid sequence of the enzyme from selected bacterial species was retrieved from KEGG database (https://www.genome.jp/kegg/) and aligned by Clustal X 2.1 (Larkin *et al*., 2007).

#### CRISPR reconstruction of each *proS* suppressor

Each *proS* suppressor mutation was reconstructed in the WT MG1655 using the scar-less CRISPR-Cas9 technology (Reisch and Prather, 2017). The specific sgRNA for each mutation was expressed from the pKDsgRNA plasmid, which was constructed by round-the-horn PCR using Phusion™ High-Fidelity DNA Polymerase (ThermoFischer Scientific) and self-ligation using T4 DNA ligase (NEB) (primers listed in Table S1). The sgRNA plasmid was introduced to the WT MG1655 strain that already contained the plasmid pCas9cr4 encoding a Cas9 protein. CRISPR-Cas9 system was induced on an LB plate containing anhydrous tetracycline (aTc) (100 ng/mL). Each *proS* mutation was assessed by PCR using allele-specific primers, and verified by Sanger sequencing. Subsequently, each strain was made into *trmD*-KO by P1 transduction without any maintenance plasmid, followed by removal of the Kan marker by FLP recombination with pCP20.

All reconstructed suppressors were confirmed for viability without the support of *trmD*. Early-log phase cultures were serially diluted, spotted on an LB plate, and grown overnight at 37 °C. To confirm the absence of TrmD from each suppressor, cell lysate proteins (15 µg) from a culture of 3 mL LB and harvested at OD = 0.4 were run on a 12% SDS-PAGE gel for Western blot analysis using an anti-TrmD primary antibody (Li and Bjork, 1999), an anti-CysRS primary antibody for a loading control, and an anti-rabbit IgG secondary antibody (Sigma-Aldrich).

#### Cellular prolyl-aminoacylation status by acid urea gel analysis

Total RNA was extracted in an acidic condition to maintain the aminoacylation status of tRNAs (Masuda *et al*., 2021). Each reconstructed *proS* suppressor lacking *trmD* was grown in LB at 37 °C to OD = 0.4, upon which cells were mixed with 10% (w/v) trichloroacetic acid (TCA) at an equal volume and incubated on ice for 10 min. Cells were harvested and resuspended in the extraction buffer (0.3 M NaOAc, pH 4.5 and 10 mM EDTA), and extracted with equal volume of ice-cold phenol-chloroform-isoamyl alcohol (25:24:1), pH 4.5. The mixtures were vortexed for 3 cycles of 1 min vortex and 1 min rest on ice, and spun at 12,000 rpm for 10 min at 4 °C. The RNA-containing aqueous phase was stored, while a second extraction of the organic phase with an equal volume of fresh extraction buffer was performed. The combined aqueous phase from the two extractions was mixed with an equal volume of cold isopropanol and incubated at −20°C for precipitation. Precipitated RNA was collected at 14,000 rpm for 20 min at 4 °C, and rinsed with wash buffer (70% ethanol and 30% 10 mM NaOAc, pH 4.5). Pellets were dried and dissolved in the buffer containing 10 mM NaOAc, pH 4.5, and 1 mM EDTA.

From each isolated total RNA sample, 10% was estimated to represent total tRNA (Masuda *et al*., 2021). Approximately 700 ng of total tRNA of each sample was mixed with a 2X loading buffer (0.1 M NaOAc, pH 5.0, 9 M Urea, 0.05% bromophenol blue (BPB), and 0.05% xylene cyanol (XC)) and separated on a 6.5%PAGE/7M urea (14 x 17 cm) in 0.1 M NaOAc, pH 5.0 at 250 V for 3 h 45 min at 4 °C. Subsequently, the gel region between XC and BPB was excised, washed in 1X Tris-borate pH 8.0 and EDTA (TBE) buffer, transferred to a wet Hybond^TM^-N+ nitrocellulose membrane (GE healthcare) in 1X TBE using Trans-Blot® TurboTM Transfer System (BIO-RAD) at constant 25 V for 20 min. The membranes were briefly air-dried, crosslinked with the bound RNA in a UV cross-linker (FB-UVXL-1000, Fisher Scientific), and then probed with a ^32^P-labeled DNA oligonucleotide targeting positions 18 to 36 of *E. coli* Pro(UGG) in a Northern blot. The membrane was imaged by a phosphorimager (Typhoon FLA 9500, GE) and bands corresponding to the charged and uncharged tRNA were quantified by ImageJ (NIH). The prolyl-aminoacylation status was calculated as the band corresponding to the charged tRNA in the sum of the bands for both charged and uncharged tRNAs.

#### Frameshift reporter assay

The nano-luciferase (nLuc) reporter gene was subcloned into plasmid pKK223-3 using NEBuilder® HiFi DNA Assembly Cloning Kit (NEB). One reporter had an insertion of the CCC triplet for Pro, while a separate reporter had an insertion of the quadruplet CCC-C codon motif (primers in Table S1). Each reporter was introduced to a reconstructed *proS* mutant expressing a normalization plasmid pZS2R-mCherry. An overnight culture in LB + Amp + Kan at 37 °C was inoculated to a fresh medium at 1:200 for WT and 1:100 for mutants. Each fresh culture was grown at 37 °C for 6 h and measured for the luciferase activity using ONE-Glo™ Luciferase Assay System (Promega) on an Infinite M200 PRO plate reader (Tecan). The luminescence at 617 nm, normalized by the mCherry fluorescence at excitation 586 nm, was used to determine the nLuc activity. The frequency of +1-frameshifting was the ratio of the CCC-C mediated nLuc activity over the CCC-mediated activity.

#### Competition fitness assay

A fitness assay was developed by mixing a reconstructed *proS* suppressor expressing the plasmid pZS2R-mCherry, and the MG1655 WT strain expressing pZS2R-YFP (Kelsic *et al*., 2015). Each strain was separately grown in LB + Kan overnight at 37 °C, and diluted 10-fold in LB. According to the OD_600_ of each, the same number of cells of each culture was inoculated to a single culture of 3 mL LB + Kan at ∼10^6^ colony-forming unit (CFU)/mL The 1:1 mixture was grown at 37 °C shaking, and an aliquot of the mixture was removed over time, diluted, and spread on an LB + Kan plate. After incubation at 37 °C overnight, the number of yellow colonies for WT and pink colonies for the mutant was counted. The plates were incubated at 4 °C as needed to enhance development of colors. The CFU of each culture at T = 0 was adjusted to 50:50, which was used to normalize the fractional distribution of the two strains over time.

#### Kinetic assay of prolyl-aminoacylation of tRNA^Pro^ *in vitro*

Each *E. coli* tRNA^Pro^ in the G37-state was synthesized by transcription, while each in the m^1^G37-state was synthesized by post-transcriptional methylation of the transcript with TrmD (Christian *et al*., 2004). Recombinant WT enzyme of *E. coli proS* was affinity-purified from *E. coli* BL21(DE3) expressing pET22b-*Ec* ProRS (Masuda *et al*., 2021). Suppressor enzymes of *proS*-D98N, *proS*-T199I, and *proS*-N232S were purified similarly from strains harboring the designated mutations but expressed in ArcticExpress (DE3) (Agilent). Each tRNA substrate was heat-denatured at 85 °C for 3 min and re-annealed at 37 °C for 15 min. Aminoacylation in steady-state conditions was performed at 37 °C in a 24 µL reaction of 0.2 to 20 µM tRNA, 0.5 to 40 nM ProRS, and 20 µM [^3^H]-Proline (Perkin Elmer, 7.5 Ci/mmol) in the aminoacylation buffer of 20 mM KCl, 10 mM MgCl_2_, 4 mM DTT, 0.15 mg/mL bovine serum albumin, 2 mM ATP (pH 8.0), and 50 mM Tris-HCl, pH 7.5. Reaction aliquots of 4 µL were removed at different time intervals and precipitated with 5% (w/v) TCA on filter pads for 10 min twice. The filter pads were washed with 95% ethanol twice, with ether once, air-dried, and measured for radioactivity in Tri-Carb 4910 TR scintillation counter (Perkin Elmer). Counts were converted to pmoles based on the specific activity of the [^3^H]-Proline after correction for signal quenching by filter pads. Data corresponding to the initial rate of aminoacylation as a function of tRNA concentration were fit to the Michaelis-Menten equation to derive the *K*_m_ (tRNA), *k*_cat_ (catalytic turnover of the enzyme), and *k*_cat_/*K*_m_ (tRNA) (the catalytic efficiency of aminoacylation) using Kaleidagraph v. 4.5 (Synergy software).

#### Evaluation of tRNA^Pro^ isoacceptors for cell viability

In *E. coli*, the Pro(GGG) and Pro(CGG) isoacceptors are non-essential and can be deleted. To test their role in cell viability, several reconstructed *proS* suppressor strains were transformed to harbor the maintenance plasmid pACYC-*araC*-P_C_-P_BAD_-human *trm5* (Masuda *et al*., 2019) to afford analysis of both the m^1^G37+ and m^1^G37− conditions. Deletion of one of these isoacceptors was made by λRed recombination (primers in Table S1) on the WT MG1655 strain to replace the respective gene with a Kan marker (*proL* for Pro(GGG) and *proK* for Pro(CGG)). P1 lysates of the *proL*-KO strain (Masuda *et al*., 2019) and of the *proK*-KO strain were used to transduce the reconstructed *proS* suppressor strains harboring the *trm5*-maintenance plasmid. After removal of the Kan marker by FLP recombination with pCP20, each deletion strain was maintained in the presence of Ara until analysis of cell viability in the m^1^G37− condition.

To test cell viability, each strain was grown overnight at 37 °C in LB + chloramphenicol (Cm) (34 µg/mL) + Ara (0.2% (w/v)). Each strain was inoculated 1:100 to an Ara-free medium of LB + Cm and grown at 37 °C for 5 h for depletion of the Trm5 protein and m^1^G37-tRNAs. Cells were then streaked out in the m^1^G37+ condition (LB + Cm + Ara) or the m^1^G37− condition (LB + Cm) and grown overnight at 37 °C.

Growth complementation analysis was achieved by introducing each isoacceptor with an over-expression pKK223-3 plasmid. The *proK* gene was synthesized by Sequenase extension and cloned into pKK223-3 (primers in Table S1). The resultant *proK*-pKK223-3 plasmid, and *proL*-pKK223-3 and *proM*-pKK223-3 plasmids (Gamper *et al*., 2015; Masuda *et al*., 2021), was separately introduced to reconstructed *proS* suppressors harboring the *trm5* maintenance plasmid. A variant of the *proM*-pKK223-3 plasmid, carrying the GGG anticodon replacing the UGG anticodon was made by site-direct mutagenesis (Masuda *et al*., 2018). A control plasmid, over-expressing Met(CAU) (initiator tRNA^Met^ encoded by *metV*) was chosen. Expression of each tRNA in MG1655 was constitutive, due to lack of the *lacI* repressor. Growth complementation was tested in both m^1^G37+ and m^1^G37− conditions at 37 °C.

#### Codon usage analysis of CC[C/U] codons in *E. coli*

The coding sequences (CDS) of *E. coli* str. K-12 substr. MG1655 were obtained from NCBI (GenBank assembly accession: GCA_000005845.2). The frequencies of each Pro codon were calculated relative to the total number of all Pro codons in each CDS using Python (version 3.7.7). GO analysis was used to predict pathways with enriched codon usage of CC[C/U]. From the 3,876 genes analyzed, those with the frequency of CC[C/U] usage equal to or higher than 0.5 (792 genes, the top 20.4%) were selected and categorized into 4 groups according to their biological processes (http://geneontology.org/).

#### Temperature-sensitivity assay

The *E. coli* MG1655 WT strain and reconstructed *proS* suppressor strains were grown in LB overnight, then spotted in serial dilution on an LB plate and grown at various temperatures (37, 30, 22 (room temp), and 18 °C) for one to three overnights. Rescue of cold sensitivity was determined by over-expression of individual tRNA^Pro^ species from the pKK223-3 plasmid.

#### Phylogenetic analysis of prolyl isoacceptors across bacterial species

Bacterial tRNA genes were retrieved from the Genomic tRNA Database (GtRNAdb) (Chan and Lowe, 2016). The number of each tRNA gene was counted for each genome. The Biopython Entrez package designed for searching genome information in NCBI was used in a custom Python script to obtain the taxonomy classifications (phylum and genus) of each genome based on the taxonomy IDs. Assemblies lacking Pro(UGG) isoacceptors for 4 box tRNAs were removed from analysis, which was likely caused by an incomplete assembly (3 assemblies). Genera with three or more representative species were selected for further analysis. Genus-level frequencies of each prolyl isoacceptor were computed by normalizing the number of species with each isoacceptor by the total number of species in the genus within this curated set. Genera were categorized into Gram-positive and Gram-negative using information obtained from PubMed literature search. The same search was used to highlight genera containing medically significant species.

To present the distribution of prolyl isoacceptors across the phylogenetic tree of bacteria, sequences of the 16S and 23S ribosomal RNA (rRNA) of each genome were retrieved from NCBI Refseq (O’Leary *et al*., 2016) or GenBank (Sayers *et al*., 2022). If rRNA annotations of a particular genome were not available, it was excluded from further analysis (3958 retrieved, 86 skipped). Sequences of 16S and 23S rRNAs were aligned using the SINA aligner (Pruesse *et al*., 2012) with the non-redundant SSU (small subunit of the ribosome) and LSU (large subunit of the ribosome) reference datasets (Ref NR 99) from SILVA (Quast *et al*., 2013) respectively. The 16S and 23S alignments of each genome were concatenated to a single FASTA entry. A single representative sequence was chosen arbitrarily from each genus selected for frequency analysis of a prolyl tRNA gene. A maximum-likelihood phylogenetic tree was generated with the prepared concatenated 16S and 23S alignments using FastTree (Price *et al*., 2009; 2010). Tree image was created using matplotlib with the ETE toolkit (Huerta-Cepas *et al*., 2016).

## SUPPLEMENTAL FIGURES

**Figure S1.**
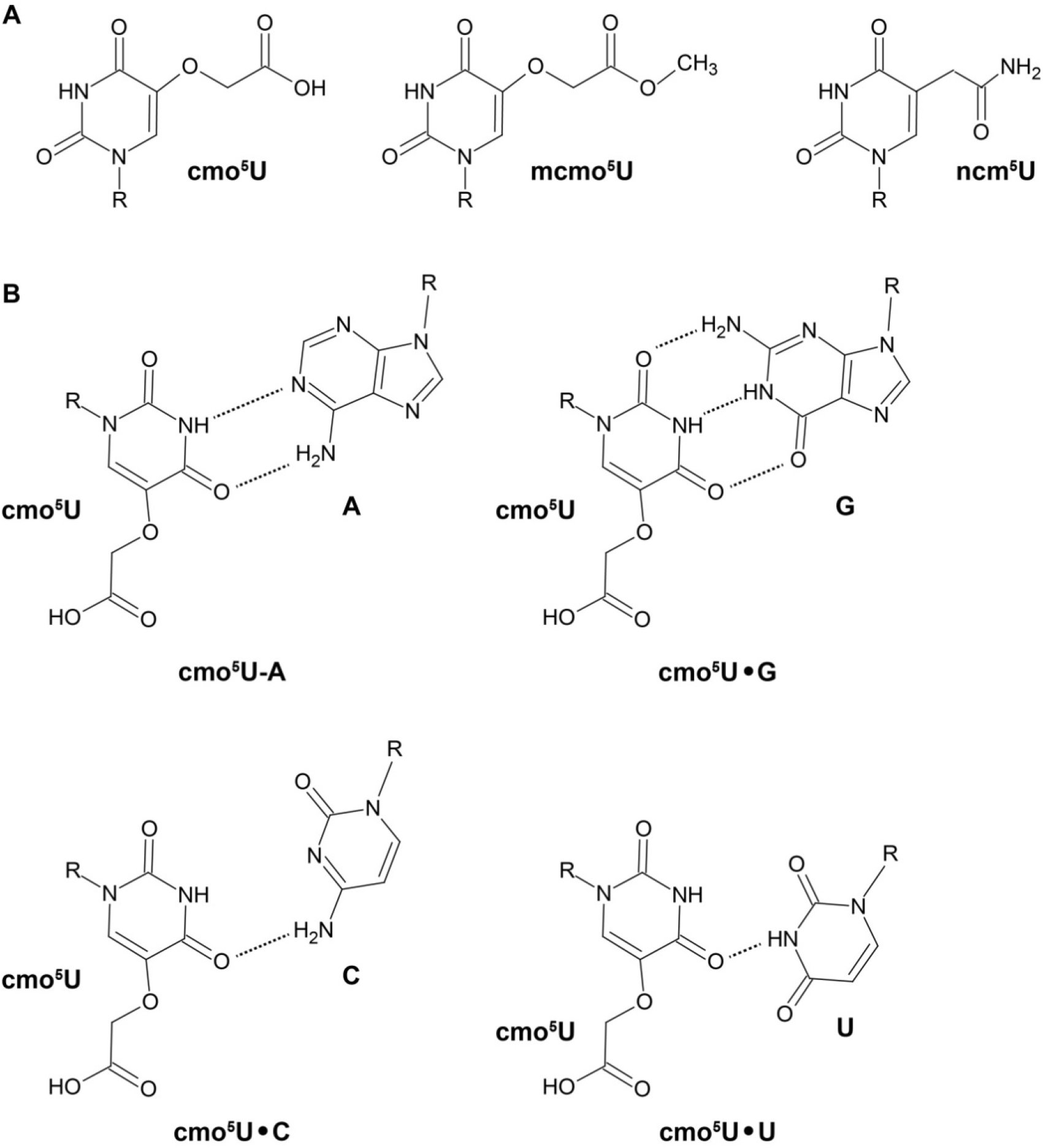
(Related to Figure 1) The cmo^5^U34 modification expands the decoding capacity using various wobble base pairing structures. **(A)** Chemical structures of cmo^5^U, mcmo^5^U, and ncm^5^U. **(B)** A model of expanded base-pairing by cmo^5^U with A, G, C, and U. The hydrogen bonds are shown by dotted lines.

**Figure S2.**
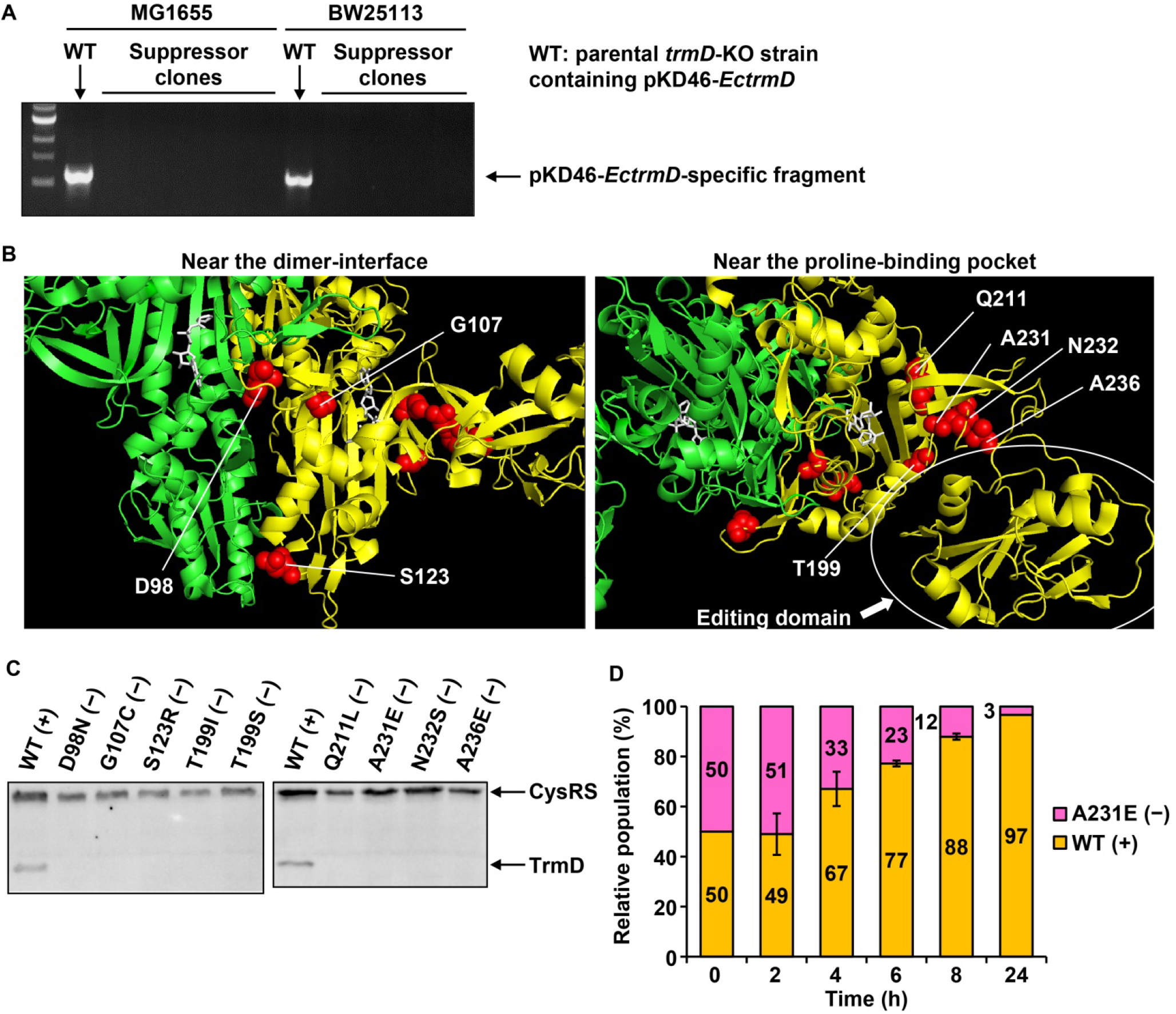
(Related to Figures 1 and 2) The isolated *proS* suppressor mutants are viable without TrmD but grow slower than the WT expressing TrmD. **(A)** Absence of the pKD46-*EctrmD* maintenance plasmid from suppressors of the *trmD*-KO allele. Suppressors were analyzed for the 5 colonies isolated from MG1655 and the 4 colonies isolated from BW25113. Absence of the plasmid was analyzed by PCR using primers specific to pKD46-*EctrmD*. The WT sample was isolated from the parental strain containing pKD46-*EctrmD*. **(B)** Mapping of *proS* suppressor mutations to the structure of *E. faecalis proS* enzyme as in Figure 1D. The images are blown up for Cluster I mutations near the monomer-monomer interphase (left) and for Clusters II and III mutations near the proline-binding pocket adjacent to the proof-reading domain (right). **(C)** Absence of TrmD in the reconstructed *proS* suppressors in Western blots. Whole-cell lysates of early-log phase cultures were run on a 10% SDS-PAGE gel. Proteins transferred to a PVDF membrane were probed by anti-TrmD and anti-CysRS antibodies. **(D)** The *proS*-A231E (−) suppressor is out-competed by the WT (+) strain in a cell fitness assay as in Figure 2C. The bar graph shows the relative abundance of the two strains in a 1:1 mixture over the time course of 24 h.

**Figure S3.**
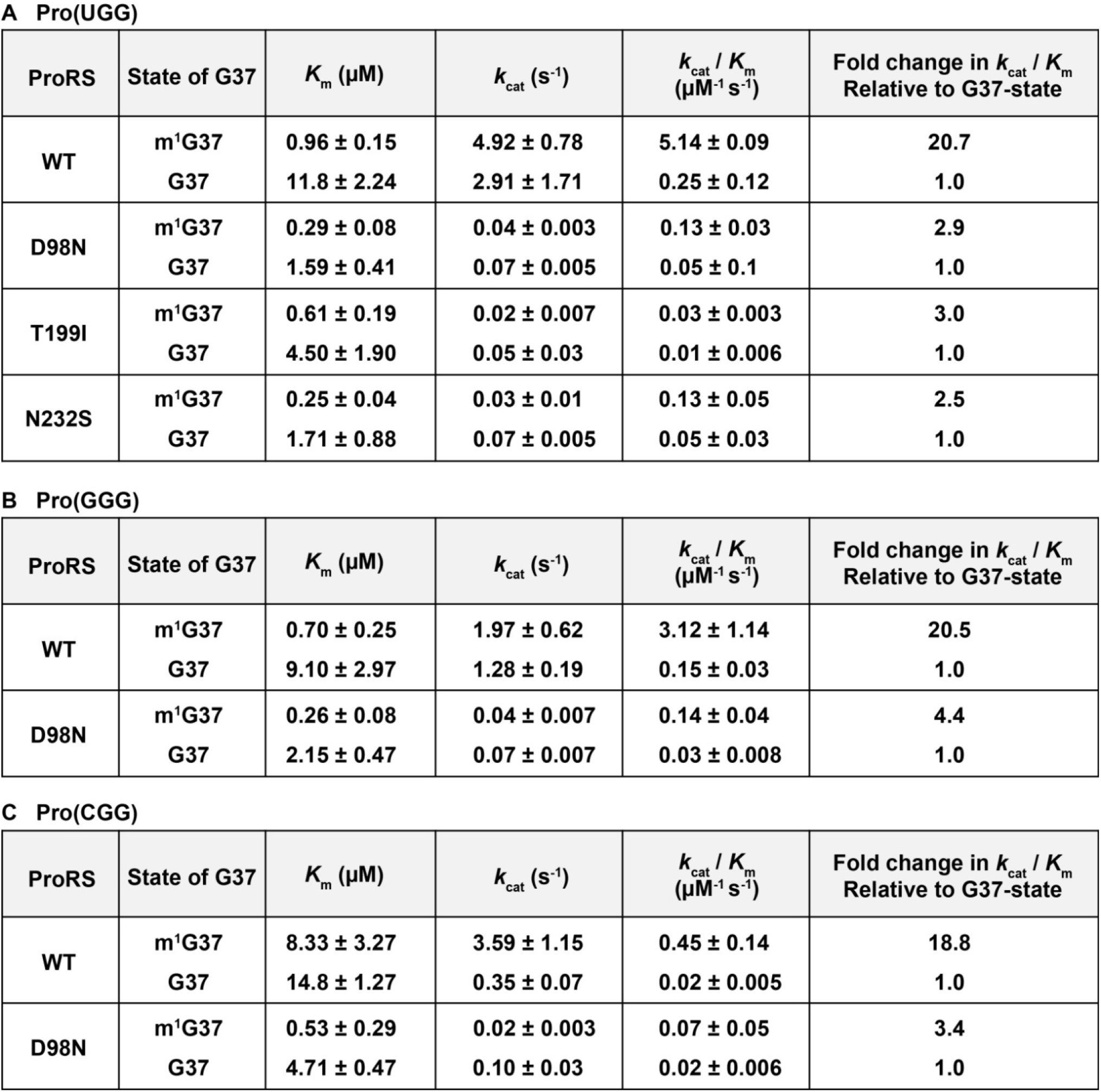
(Related to Figure 3) Kinetic parameters of prolyl-aminoacylation. **(A)** Kinetic parameters of prolyl-aminoacylation of Pro(UGG) shown in Figure 3B. For each enzyme, *K*_m_ (µM) for the tRNA substrate, *k*_cat_ (s^-1^), *k*_cat_/*K*_m_ (µM^-1^s^-1^), and the fold-change of *k*_cat_/*K*_m_ of the G37-state relative to the m^1^G37-state are shown. **(B)** Kinetic parameters of prolyl-aminoacylation of Pro(GGG) shown in Figure 3C. **(C)** Kinetic parameters of prolyl-aminoacylation of Pro(CGG) shown in Figure 3D.

**Figure S4.**
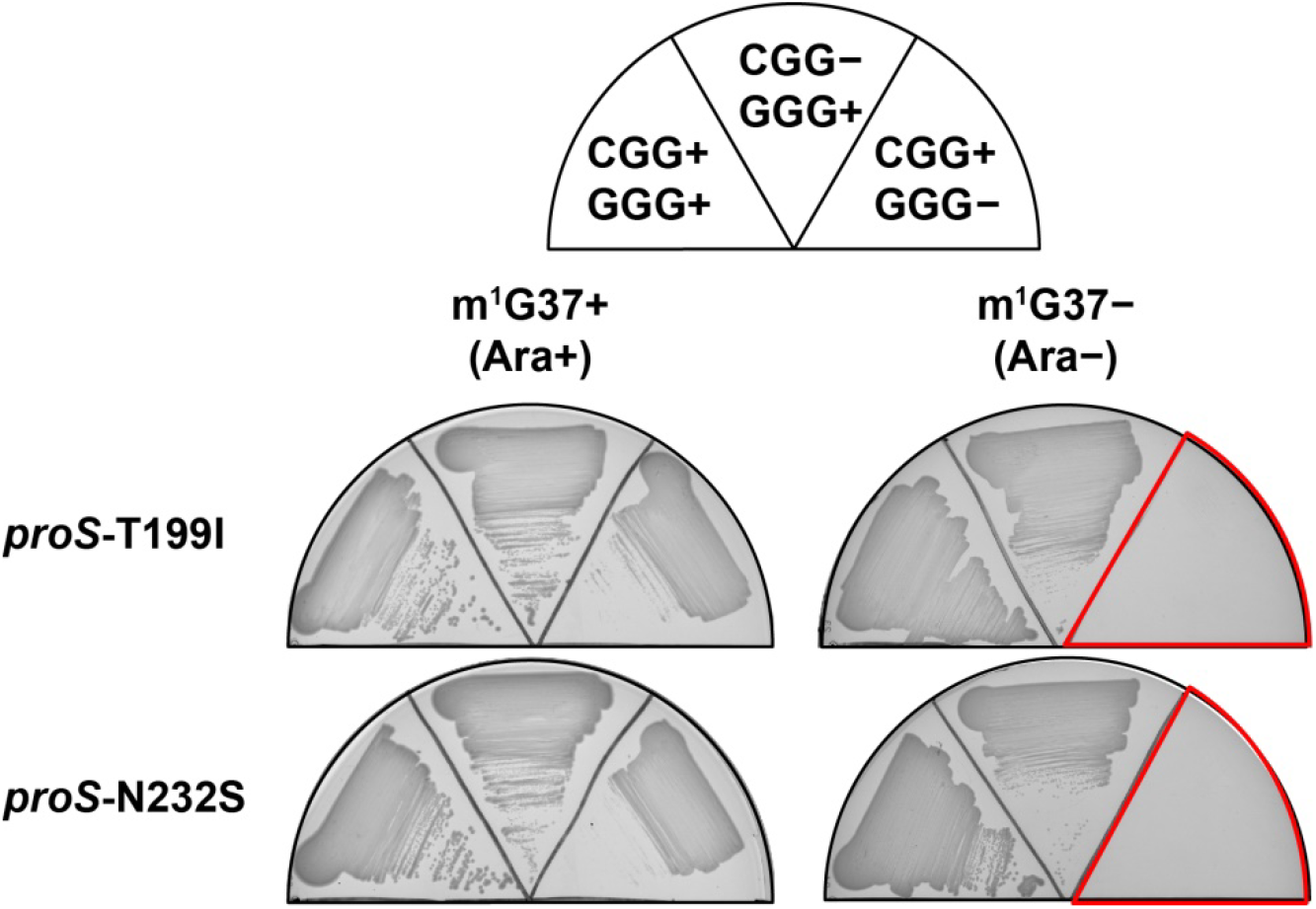
(Related to Figure 4) Requirement of Pro(GGG) for cell viability of *proS*-T199I and *proS*-N232S mutants in m^1^G37− condition. Pro(CGG) or Pro(GGG) was removed from *proS*-T199I and *proS*-N232S mutants lacking *trmD* and cells were grown in m^1^G37+ or m^1^G37− condition as in Figure 4C. No growth was observed when both m^1^G37 and Pro(GGG) were absent.

**Figure S5.**
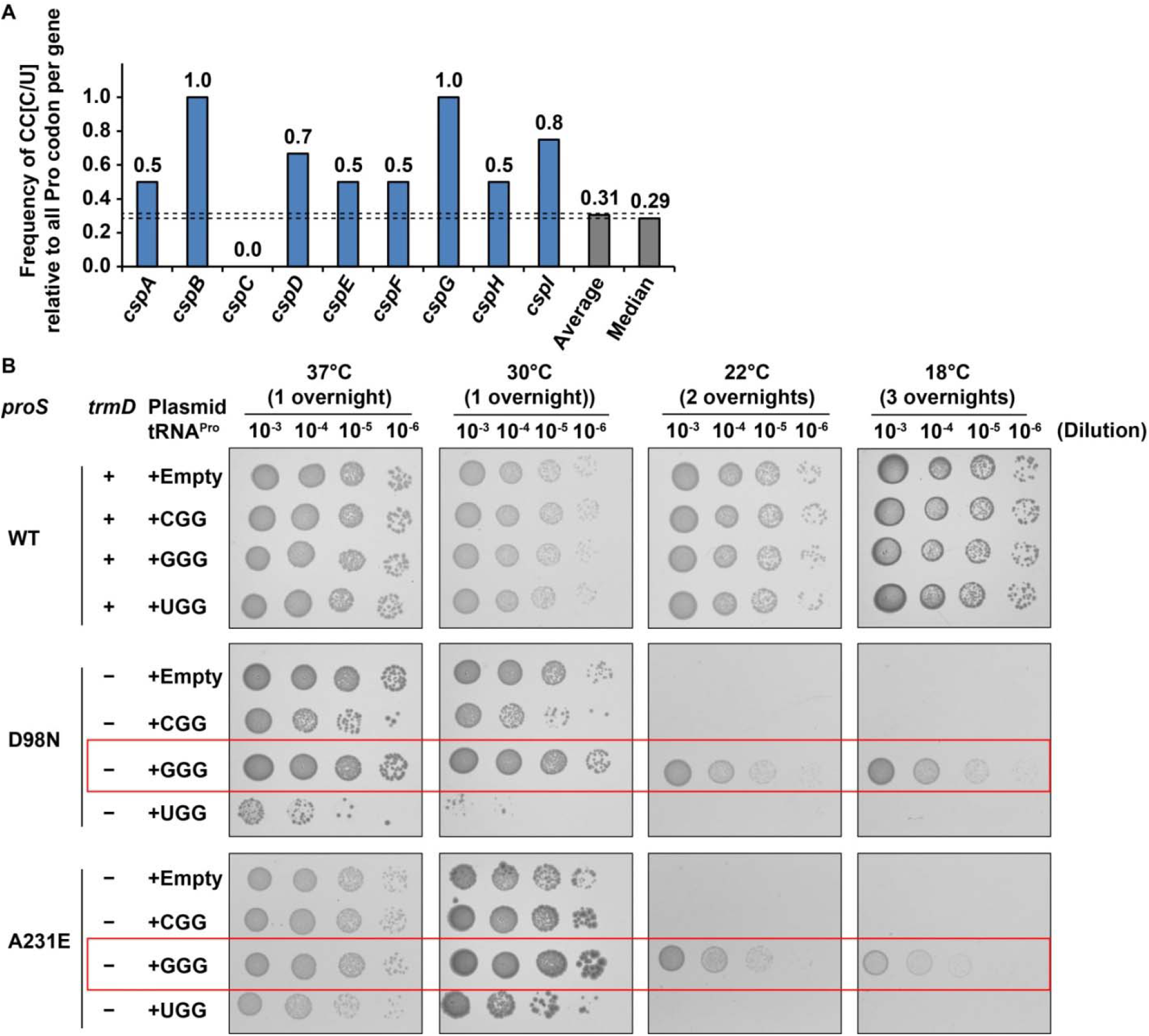
(Related to Figure 5) High frequency of CC[C/U] codons in *E. coli csp* genes. **(A)** The frequency of CC[C/U] codons is shown for selected *csp* genes, showing higher frequencies than the average and median (0.31 and 0.29, respectively) for most of these genes. **(B)** Rescue of cold sensitivity by expression of Pro(GGG) in *proS*-D98N and *proS*-A231E suppressors as in Figure 5D. The *proS*-A231E mutant was incubated at 30 °C for 2 overnights due to a slower growth phenotype.

**Figure S6.**
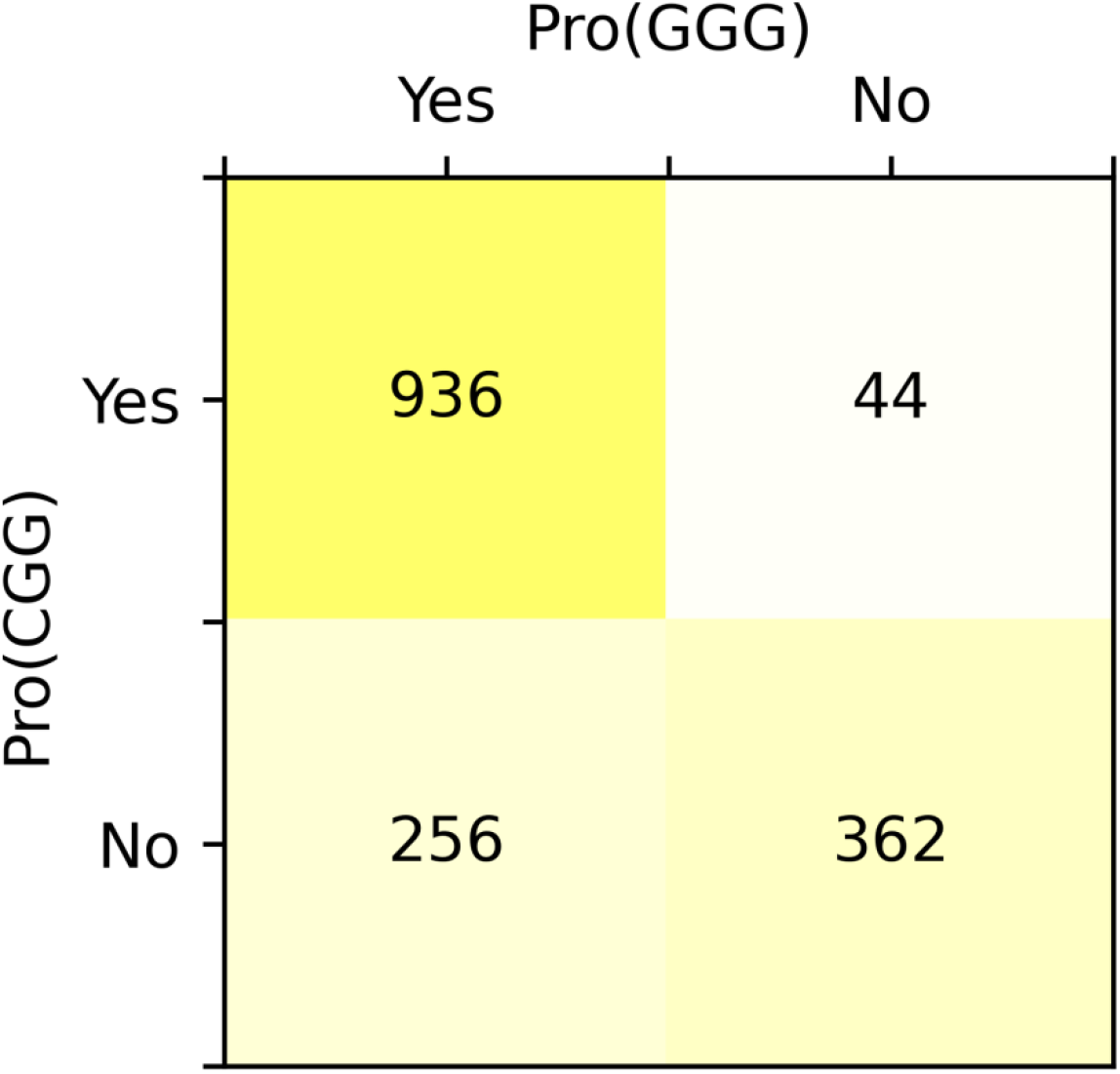
(Related to Figure 6) Co-occurrence of Pro(GGG) and Pro(CGG) across bacterial species. Heat-map table showing frequencies of co-occurrence of Pro(GGG) and Pro(CGG) across all bacterial species that are available in GtRNAdb. Counts are species within GtRNAdb beyond the curated set used in Figure 6A. Few bacterial species have Pro(CGG) but lack Pro(GGG), whereas many have Pro(GGG) but lack Pro(CGG).

## SUPPLEMENTAL TABLES

**Table S1.**
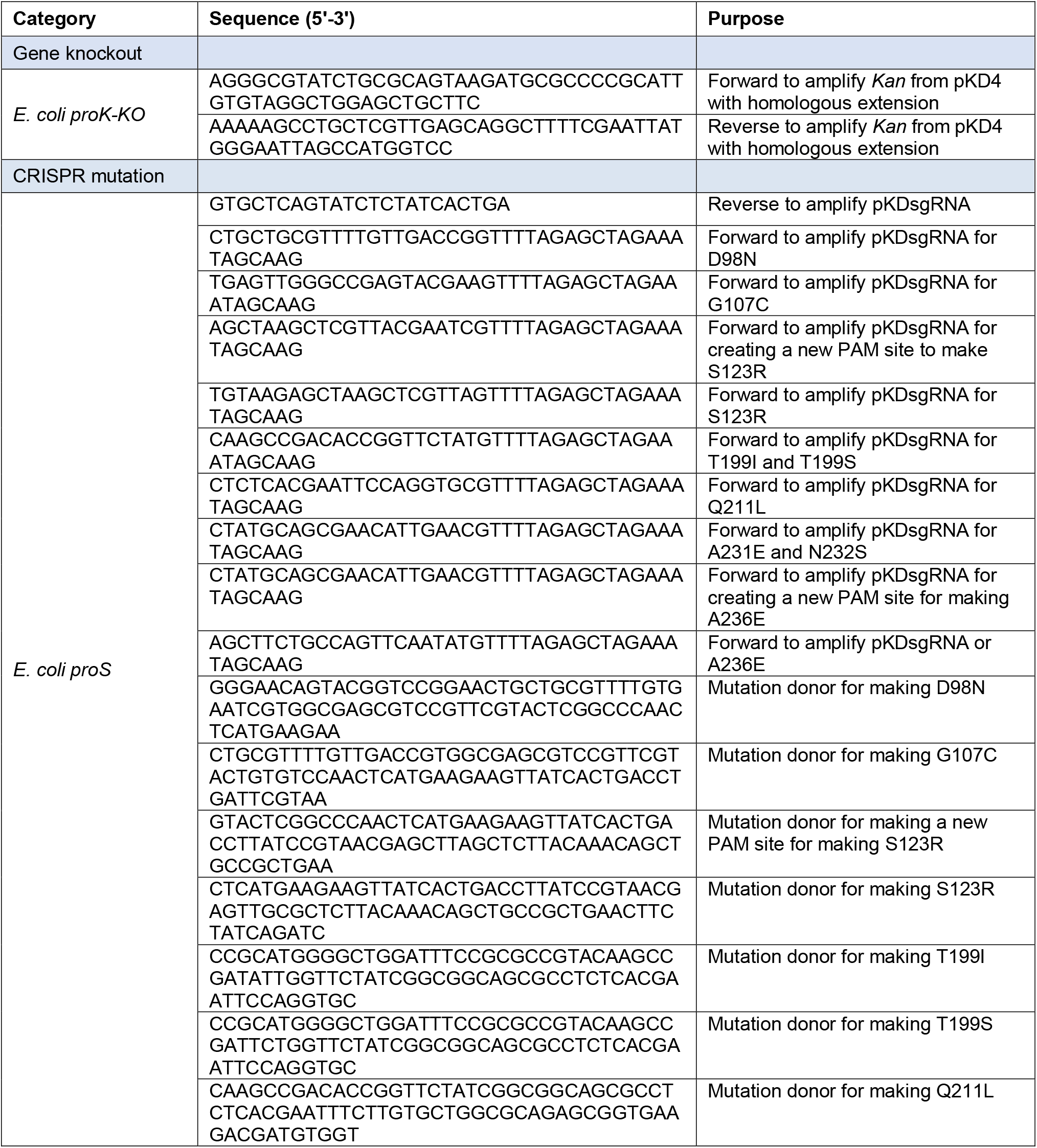
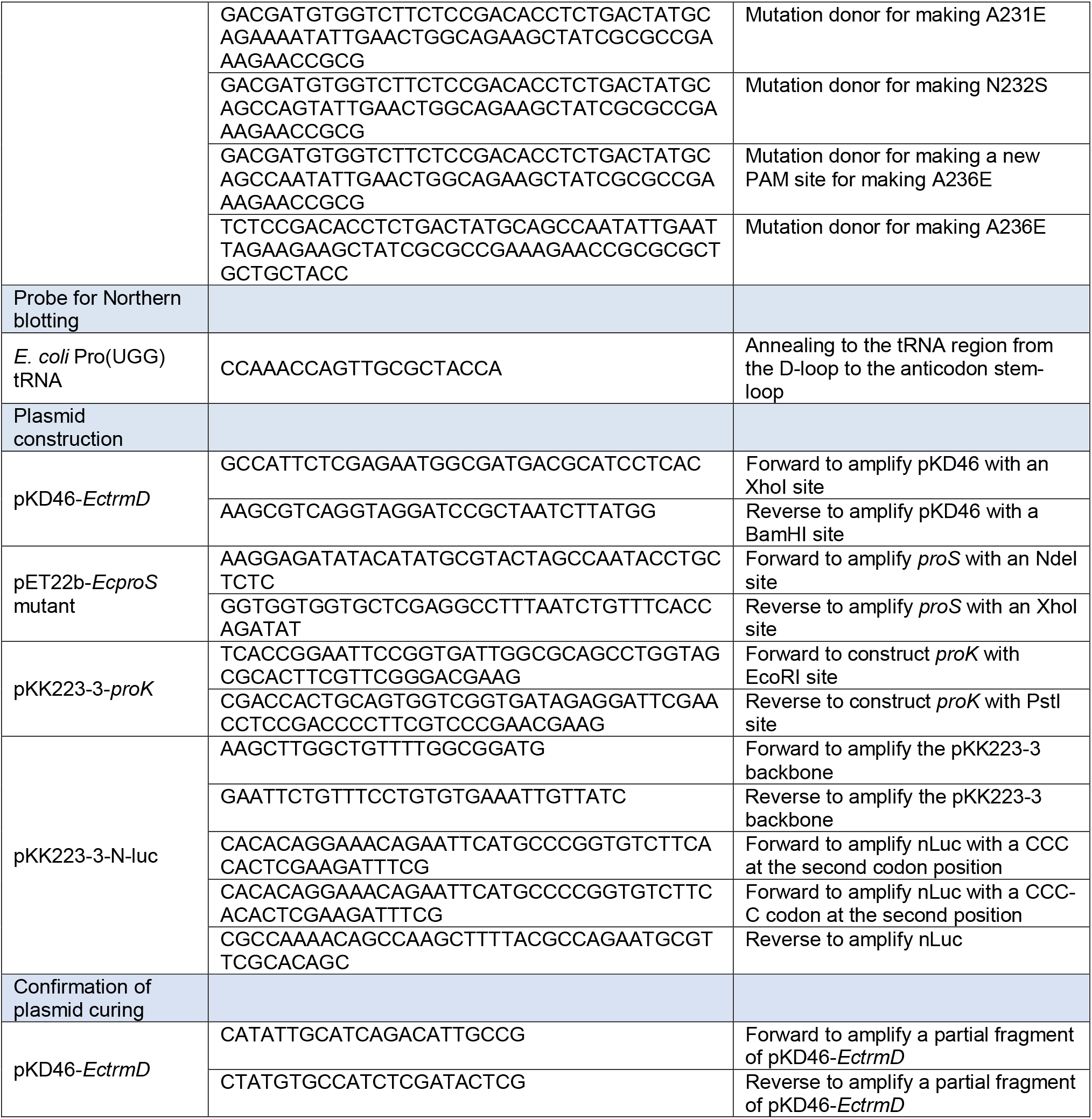
Primers used in this study.

**Table S2.**
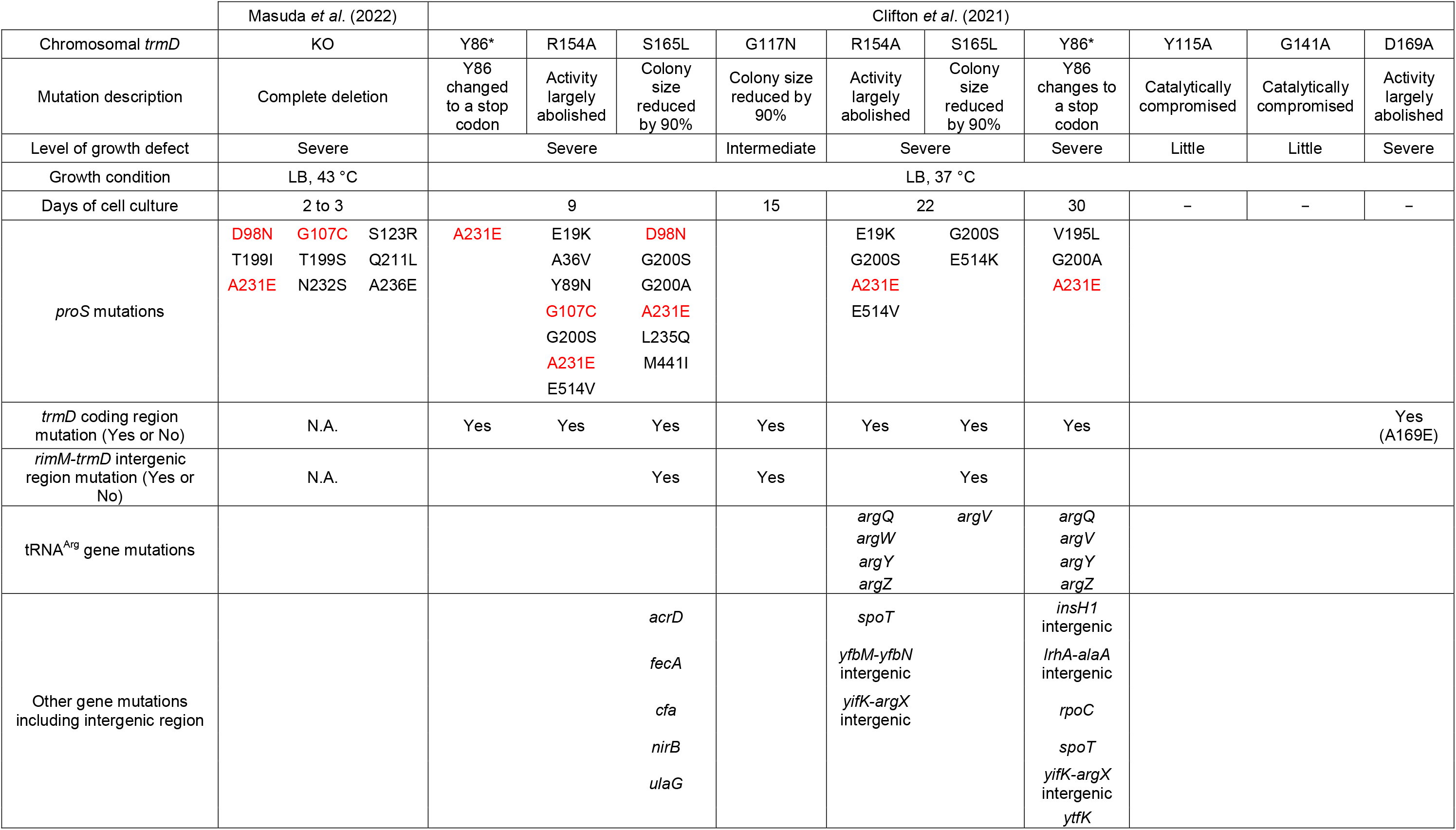
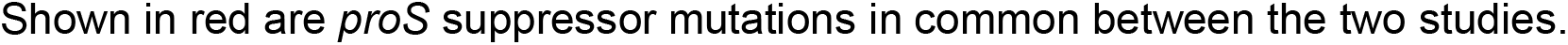
Comparison between this study by Masuda *et al*. and the study by Clifton *et al* (Clifton *et al*., 2021).

## Notes

### Competing Interest Statement

The authors have declared no competing interest.

